# SIRT5 is the desuccinylase of LDHA as novel cancer metastatic stimulator in aggressive prostate cancer

**DOI:** 10.1101/2021.08.08.455585

**Authors:** Oh Kwang Kwon, In Hyuk Bang, So Young Choi, Ju Mi Jeon, Ann-Yae Na, Yan Gao, Sam Seok Cho, Sung Hwan Ki, Youngshik Choe, Jun Nyung Lee, Yun-Sok Ha, Eun Ju Bae, Tae Gyun Kwon, Byung-Hyun Park, Sangkyu Lee

**Author notes:** Corresponding author. (Lee S), (Park B-H), (Kwon TG). Equal contribution (Oh Kwang Kwon, In Hyuk Bang, So Young Choi).

## Abstract

Prostate cancer (PCa) is the most commonly diagnosed genital cancer in men worldwide. Among patients who developed advanced PCa, 80% suffered from bone metastasis, with a sharp drop in the survival rate. Despite great efforts, the details of the mechanisms underlying castration-resistant PCa (CRPC) remains unclear. SIRT5, an NAD^+^-dependent desuccinylase, is hypothesized to be a key regulator of various cancers. However, compared to other SIRTs, the role of SIRT5 in cancer has not been extensively studied. Here, we showed significantly decreased SIRT5 levels in aggressive PCa cells relative to the PCa stages. The correlation between the decrease in the SIRT5 level and the patient’s survival rate was also confirmed. Using quantitative global succinylome analysis, we characterized a significant increase of lysine 118 succinylation (K118su) of lactate dehydrogenase A (LDHA), which plays a role in increasing LDH activity. As a substrate of SIRT5, LDHA-K118su significantly increased the migration and invasion of PCa cells and LDH activity in PCa patients. This study investigated the reduction of SIRT5 and LDHA-K118su as a novel mechanism involved in PCa progression. It can also be proposed as a new target that can prevent castration-resistant PCa progression, which is key to PCa treatment.

## Introduction

Prostate cancer (PCa) is the most common genital cancer in men, with about 164,690 new cases of PCa in 2017, as estimated by the American Cancer Society [1]. PCa was initially considered a cancer of the elderly, but PCa incidence in patients below age 55 is currently increasing by more than 10% in the United States [2]. Although the 5-year survival rate of localized PCa is > 99%, the outlook is relatively poor once PCa advances [3, 4]. Half of the men with castration-resistant PCa (CRPC) developed bone metastasis within two years of CRPC diagnosis [5]. PCa can spread to a variety of tissues, especially the lymph nodes and bone [6]. In a previous study, PCa first metastasized to the bone and then secondarily to other organs [7]. The progression from the bone to other metastatic sites is associated with a decreased survival in CRPC.

A diversity of covalent post-translational modifications (PTMs) affect various significant biological mechanisms during cancer progression [8]. Lysine residues have various PTMs, which are acylations, such as acetylation, succinylation, malonylation, and glutarylation. These acylations are regulated by silent information regulator 2-like protein (sirtuin; SIRT) while removing the modified acyl groups from proteins as deacylase. SIRTs are highly conserved proteins with NAD^+^-dependent deacylase activity and classified as class III histone deacetylases [9, 10]. There are seven SIRTs (SIRT1–SIRT7) in mammals, all of which show diverse subcellular localizations and distinct functions [11]. Numerous studies have revealed that SIRTs play critical roles in cancer progression and metastases by regulating angiogenesis, inflammation, and epithelial-to-mesenchymal transition (EMT) [12-14].

In this study, we studied the regulating factors involved when PCa secondarily metastasizes to other organs from the bone using two PCa-cell lines (PC-3 and PC-3M, a highly metastatic subline of PC-3 isolated from a PC-3-induced mouse tumor) [15]. The PC-3 cell line is commonly used to study PCa as the leading cause of death in PCa patients is bone metastasis [6, 16]. Furthermore, the PC-3M cell line used as positive control is a subline of PC-3 isolated from a PC-3-induced mouse tumor and showed increased migration compared to PC-3 cells [15].

We studied SIRT expression between two types of cell lines and found that SIRT5 expression significantly decreased with PCa progression. SIRT5 is known as the deacylase of several acylations but plays the most important role in desuccinylation [11, 17]. Various roles of lysine succinylation (Ksu) in individual proteins have been reported, particularly in relation to cancer progression and metastasis [18-20]. For example, S100A10 was found to be succinylated at lysine residue 47 in gastric cancer. SIRT5 regulates desuccinylation, and levels of succinylated S100A10 increased in human gastric cancer [21]. Thus, this study identified the specific substrate protein of SIRT5 and suggested the role of Ksu of the identified substrate protein in increasing the metastatic potential of PCa based on comparative global succinylome study in PC-3 and PC-3M cell lines.

## Results

### SIRT5 is significantly down-regulated in advanced PCa

To identify the specific SIRT isoform according to PCa progression, we determined the SIRT1 expression level through SIRT7 in four representative androgen-dependent (LNCaP and LNCaP-LN3) and androgen-independent (PC-3 and PC-3M) PCa cell lines (**Figure 1A**). As expected from previous studies, SIRT1 and SIRT7 were up-regulated in PC-3M cells^26,27^. Meanwhile, SIRT5 decreased depending on PCa progression from LNCaP to PC-3M cells. SIRT5 expression decreased in the progressed cells (LNCaP-LN3) compared to LNCaP cells and was significantly down-regulated in PC-3M as the more progressive cells than PC-3 cells. Because PC-3M exhibits enhanced metastasis compared to PC-3, decreased SIRT5 expression in PC-3M may be associated with metastasis [15]. SIRT5 is a protein in the cytosol but is primarily present in the mitochondria [11]. In the present study, we also confirmed SIRT5 distribution by evaluating confocal images and immunoblots after subcellular fractionation (**Figs. S1A and S1B**). SIRT5 in PC-3M cells was significantly down-regulated in the cytosol and the mitochondria compared to PC-3 cells. *Sirt5* mRNA expression showed no significant difference between the two cell lines due to genetic factors (**Figure S1C**). As PC-3M is a cell isolated from PC-3, there is no genetic difference regarding SIRT5 expression. It can be assumed that there is a difference in the expression of SIRT5 protein due to causes other than gene expression. In contrast to the decrease in SIRT5 expression, SIRT2 increased in aggressive cancer cells (**Figure 1A**). SIRT2 is known to increase migration and invasion in various cancers, and this study confirmed that it is increased in the aggressive PCa cell line [22, 23].

**Figure 1.**
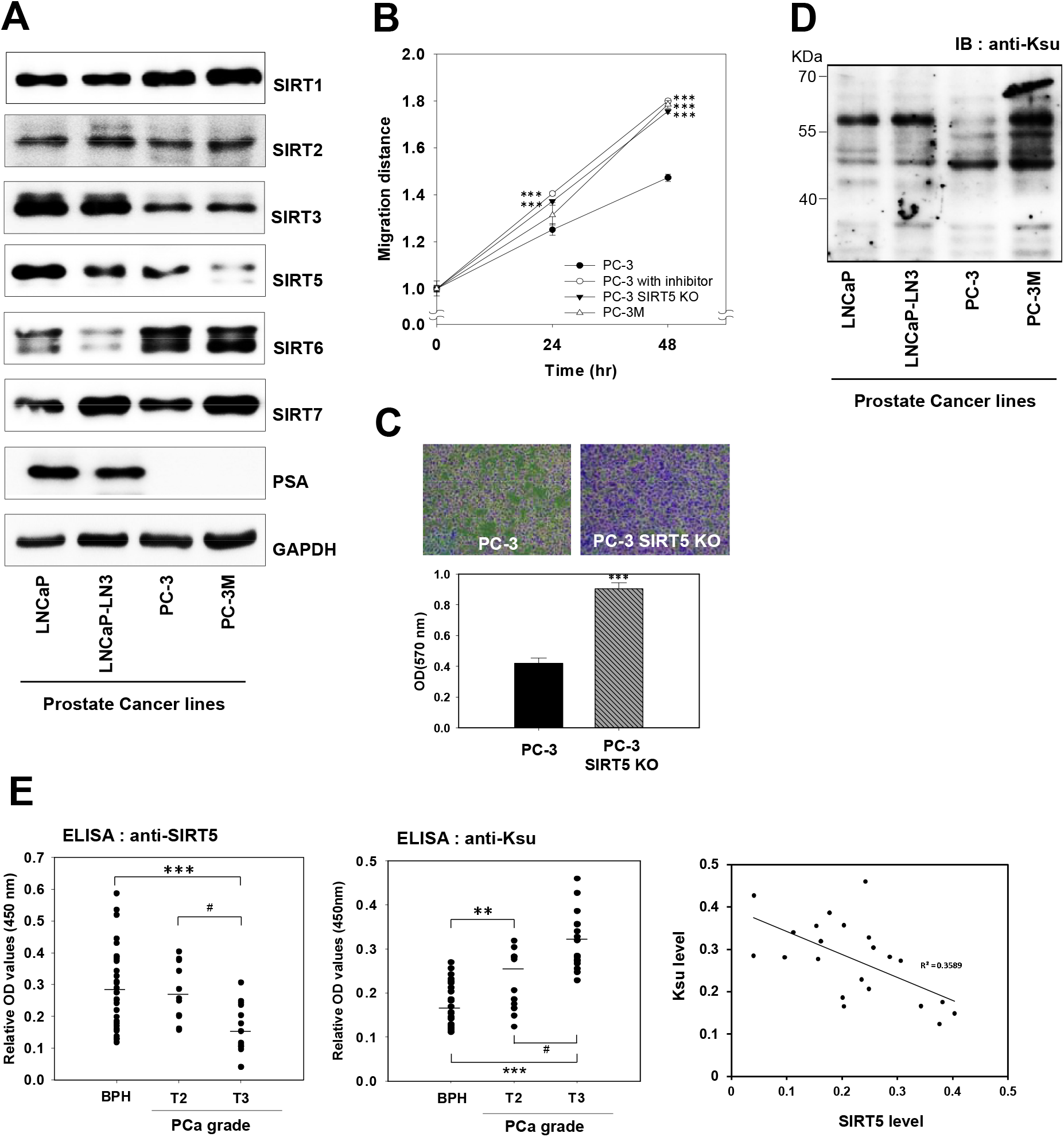
Characterization of SIRT5 in PCa. **A**. SIRT5 was significantly decreased in PC-3M compared to PC-3 cells. Levels of seven SIRT members were analyzed by Western blot with indicated antibodies in PCa cell lines. **B**. SIRT5 regulates cell migration. SIRT5 selective inhibitor (HY-112634) and SIRT5-KO promoted cell migration (n=3). Cell migration was evaluated by a wound-healing assay in PC-3 cells after treatment with an SIRT5 inhibitor and in PC-3 SIRT5-KO and PC-3M cells. **C**. *SIRT5* knockdown promoted cell invasion. Cells that invaded through Matrigel were stained with crystal violet (n=3). Representative images of invaded cells (40×). **D**. The global Ksu levels increased in the PC-3M cells. Levels of Ksu were analyzed using Western blot with anti-pan-Ksu antibody in whole lysate-derived PCa cells. **E**. SIRT5 was significantly decreased depending on the stage of PCa. In contrast, Ksu was significantly increased depending on the stage of PCa. The coefficient between SIRT5 expression and Ksu levels was determined in the scatter plot in PCa tissues. Levels of SIRT5 and Ksu were confirmed based on sandwich ELISA in BPH, T2, and T3 grade PCa tissues with indicated antibody. Data are mean ± SE. **p < 0.01, ***p < 0.001 vs. BPH; #p < 0.01 vs. T2. See also **Figures S1** and **S2**.

### SIRT5 significantly reduces cell migration and invasion in PCa

To test the role of SIRT5 in PCa cells, whether SIRT5 affects the migration and invasion of PCa cells was assessed. Migration assays were performed using a selective peptide inhibitor of SIRT5 to determine whether SIRT5 regulates PCa cell migratory ability [24][25] (**Figures 1B** and **S1D**). The results showed that migration ability was increased when SIRT5 activity was inhibited. Next, the clustered regularly interspaced short palindromic repeats (CRISPR)/CRISPR-associated protein-9 endonuclease (Cas9) genome editing system was used to knock out the *SIRT5* gene in PC-3 cells (PC-3 SIRT5-KO). When PC-3 SIRT5-KO cells were evaluated for selective KO efficiency by Western blot, SIRT5 levels were markedly decreased, and no changes were observed in the levels of other SIRT proteins (**Figure S1E**). To assess whether SIRT5 affects cellular growth rate, the Cell Counting Kit-8 (CCK-8) assay [26, 27] was used to compare the proliferation of normal (PC-3), control (vehicle), PC-3 SIRT5-KO, and PC-3M (positive control) cells (**Figure S1F**). This assay revealed that SIRT5-KO cells proliferated significantly faster than normal PC-3 cells (p < 0.001). This result indicates that SIRT5 can regulate the proliferation of the PC-3 cell line.

Next, a migration assay [27, 28] with PC-3, PC-3 SIRT5-KO, and PC-3M cells was used to assess whether the migratory ability is dependent on SIRT5. Interestingly, SIRT5-KO cells migrated significantly faster than PC-3 cells as measured by TScratch software, suggesting that SIRT5 regulates the migration of PCa cells, supporting the increase in migration when treated with SIRT5 inhibitors (**Figures 1B and S1D**). Furthermore, invasion assays [29] were performed to evaluate the invasiveness of PCa cells upon SIRT5-KO (**Figure 1C**). The results demonstrated an increased invasion of PC-3 SIRT5-KO cells through the Matrigel to the bottom membrane. These two cell lines were eluted with methanol and measured at 570 nm [29]. Invasion assay data for PC-3 and PC-3 SIRT5-KO demonstrated that SIRT5 depletion in PC-3 cells is associated with increased invasiveness. These results indicate that SIRT5 is an important factor in increasing cell migration related to PCa progression.

### Decreased SIRT5 is associated with PCa progression

In **Figure 1A**, SIRT5 acting as a representative desuccinylase was decreased in PC-3M cells of the highly metastatic PCa cell line. Due to the decrease in the level of SIRT5, it was observed that the Ksu level increased in PC-3M cells (**Figure 1D**). The increased Ksu level additionally showed the most change in expression among other acylations, including K-acetylation, K-β-hydroxybutylation, K-malonylation, and K-glutarylation in metastatic PCa cell lines (**Figure S1G**). Furthermore, low levels of SIRT5 were significantly correlated with T grade increase (T2 to T3) but less strongly associβatedβ βwith age, weight, body mass index, and Gleason score (**Table S1**). To verify the correlation between SIRT5 and PCa progression, an enzyme-linked immunosorbent assay (ELISA) was carried out to assess SIRT5 and Ksu levels in PCa tissues [benign prostatic hyperplasia (BPH) as control and T2 and T3 stages; **Figure 1E**]. ELISA showed that SIRT5 levels were significantly decreased in more advanced PCa tissues, as SIRT5 was reduced in PC-3M, which are more highly metastatic PCa cells. Corresponding to the decreased SIRT5 level in PCa tissues, the Ksu level in T3 patients compared to T2 patients’ tissue was significantly increased, representing a result of reduced desuccinylation activity. Moreover, SIRT5 was negatively correlated with Ksu in patients’ tissue (**Figure 1E**). Patients with a lower SIRT5 level showed a worse overall survival, although it was not significant (**Figure S2**). Although the results were not significant due to the lack of sufficient observation time and the number of patients due to the long progression of PCa, a correlation between reduced survival rate and decreased SIRT5 was observed.

### Succinylated proteins are identified by global succinylome research

Based on the results, it was hypothesized that the decrease of SIRT5 level promotes the PCa progression. To explore the substrate of SIRT5 associated with aggressive PCa, a global protein succinylation assay was performed in PCa cell lines using SILAC-based proteomics and global immunoprecipitation (IP) enrichment [30] (**Figure 2A**). PC-3 and PC-3M cell lines were labeled with light (K0R0) and heavy (K8R10) SILAC medium, respectively. Biological replicates of two different samples were also prepared to minimize biological variability. The Pearson correlation coefficients of the ratio of proteins and Ksu peptides were 0.89 and 0.71, respectively (**Figure S3A**). Ksu analysis showed that 81 of 169 (47.9%) of Ksu proteins were overlapping and 167 of the 488 (37.3%) Ksu sites were duplicated.

**Figure 2.**
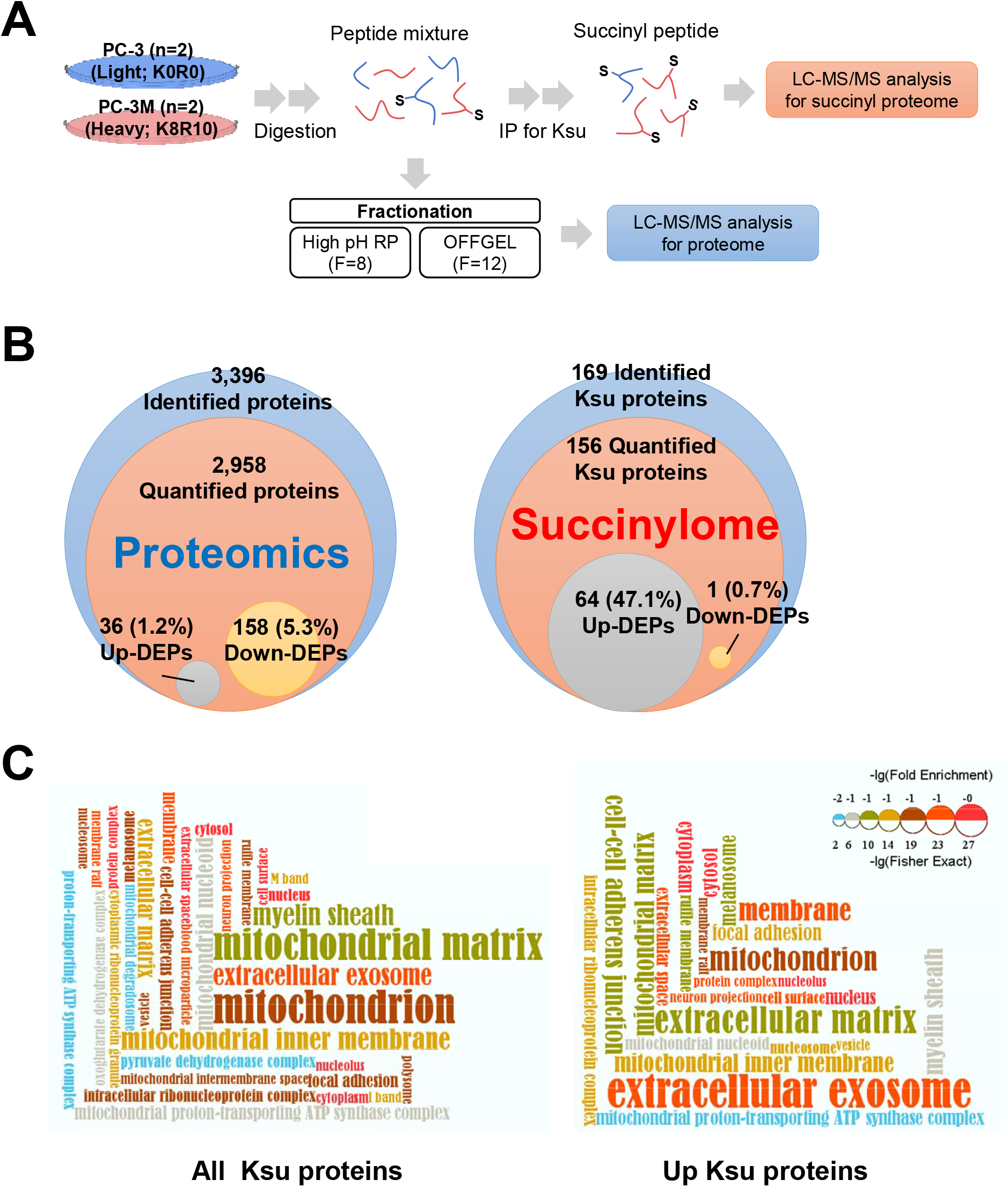
Increased Ksu in the cytoplasm in PCa. **A**. Change in Ksu between PC-3 and PC-3M cells was identified by the global lysine succinylome technique. PC-3 and PC-3M cells were labeled with light or heavy amino acids in SILAC media, respectively. The peptide mixtures were enriched by IP using anti-succinyl-lysine antibody-conjugated agarose beads (n = 2). The succinyl-lysine proteins were identified and quantified by high-resolution tandem MS (MS/MS) analysis and peak alignment. **B**. The succinylation level is changed more than protein changes. Only 6.3% of the total protein was DEPs, but 48.8% of succinylated proteins were changed. **C**. The succinylation level of the cytoplasm and extracellular components is further increased than mitochondria. All Ksu and up-regulated Ksu proteins in PC-3M cells were visualized by functional enrichment of the GO Cell Component category. See also **Figure S3** and **Tables S2** to **S4**.

This study was able to quantify 2958 proteins and 194 (6.5%) differentially expressed proteins (DEPs), which included 36 (1.2%) up-regulated and 158 (5.3%) down-regulated proteins in PC-3M cells compared to PC-3 cells (**Figure 2B**; **Table S2**). From the succinylome data, 448 Ksu sites on 169 proteins were identified, and 406 of these Ksu sites in 136 proteins were quantified and normalized to protein levels (**Table S3**). Interestingly, 144 of 145 differentially expressed sites of Ksu were found to be increased in PC-3M cells compared to PC-3 cells. This accounted for 35.5% of the total quantified Ksu sites and 47.1% of the quantified Ksu proteins (**Figure 2B**). As for the proteins, the difference between the two cell lines is insignificant as only 6.5% of the cells are DEP, whereas in the case of Ksu, 47% were DEP. Therefore, it is suggested that protein succinylation is a more important factor for the aggressiveness of PCa cells.Gene Ontology (GO) enrichment analysis was conducted using DAVID to compare the biological properties of total proteins containing Ksu (Ksu protein) and up-regulated Ksu proteins (**Figure S3B**) [31]. The DAVID analysis result suggested that different categories were enriched between identified total Ksu proteins and up-regulated Ksu proteins in the GO Biological Process (GOBP), GO Cell Component (GOCC), and GO Molecular Function (GOMF). Furthermore, in the GOCC results, the subcellular distribution results of total Ksu protein, and up-regulated Ksu protein were also different (**Figure 2C**). Intriguingly, total Ksu proteins appeared to be more abundant in the mitochondria (42.9%) than in the cytoplasm or nucleus. In contrast, Ksu proteins up-regulated in PC-3M cells were more dominant in the cytoplasm (46.3%). Therefore, it could be suggested that the substrate of SIRT5, which affects PCa progression, is a cytosol protein.

According to the above results of the bioinformatic analysis of the succinylome in advanced PCa, Ksu proteins up-regulated in PC-3M cells were dominated by extracellular and cytoplasmic proteins but not by mitochondrial proteins (**Figure 2C**). Therefore, among the extracellular exosome and extracellular matrix proteins in the GOCC category, 67 up-regulated Ksu sites on 26 proteins, excluding mitochondrial, mitochondrial matrix, and mitochondrial inner membrane proteins, were selected. Furthermore, 36 increased Ksu sites in 15 cytoplasmic proteins based on the subcellular location results were selected (**Figure S3C**). Finally, 33 up-regulated Ksu sites on 12 proteins, characteristically extracellular or cytoplasmic while excluding the mitochondrion, as candidate substrates for aggressive PCa were selected (**Figure S3D**; **Table S4**). Of the 12 proteins, lactate dehydrogenase A (LDHA) was selected because four of the seven sites capable of succinylation were actually succinylated, and LDHA is known as a key player related to carcinogenesis in many previous studies [32, 33].

### LDHA-K118 succinylation (K118su) increases LDH activity

It could be estimated that the high migration and invasion of PC-3M cells are related to LDHA succinylation as a substrate of SIRT5. LDH activity was significantly increased in SIRT5-KO and PC-3M cells compared to PC-3 cells. However, there was no change in LDHA expression in the four cells evaluated (**Figures 3A and S4A**). LDHA expression in wild-type (WT) and SIRT5-KO cell lines of PC-3 cells was not changed, but succinylation was increased in SIRT5-KO cells (**Figure S4B**). Also, K-acetylation, which promotes LDHA degradation, was reduced in SIRT5-KO cells (**Figure S4B**) [22]. LDH activity also increased in a concentration-dependent manner, even when PC-3 cells were treated with an SIRT5 selective inhibitor (HY-112634) (**Figure 3B**). Likewise, there was no change in LDHA according to treatment with SIRT5 inhibitor, but LDHA succinylation increased in the same pattern as activity increased (**Figure S4C**). To verify whether LDHA succinylation increases LDH activity, succinylation of LDHA was induced by treatment with succinate in PC-3 cells. Succinate treatment in PC-3 cells increased the succinylation of LDHA, and also significantly increased the activity of LDH (**Figures 3C and S4D**). Additionally, cell migration and proliferation were increased by succinate treatment (**Figure S5**). As a result, the increased succinylation in LDHA linked to increase of LDH activity, cell migration and proliferation.

**Figure 3.**
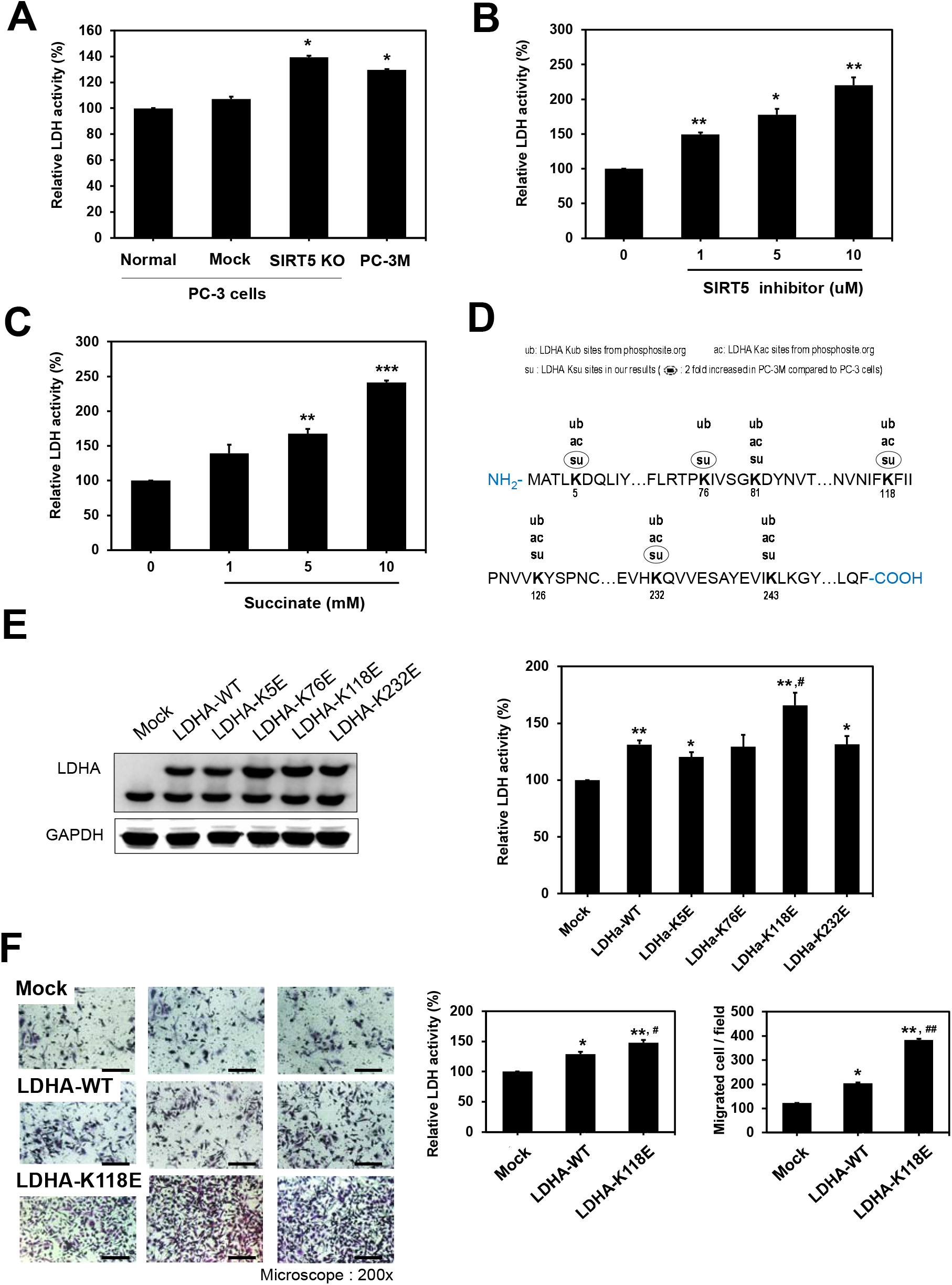
LDHA desuccinylation by SIRT5 is related to aggressive PCa. **A**. LDHA activity was increased in SIRT5-depleted PCa, PC-3M, and PC-3 SIRT5-KO cells. **B**. SIRT5 inhibition was increased by LDHA activity. LDHA activity was determined by the treatment with the SIRT5 selective inhibitor (HY-112634) in PC-3 cells. **C**. Inducing succinylation increased LDHA activity. Succinate was administered to PC-3 cells to increase the succinylation level, and LDHA activity was determined. **D**. Detected succinylated sites in LDHA. Among the detected sites, sites that increased more than two times in PC-3M were marked with circles. The acetylation and ubiquitylation sites known in previous studies are indicated. **E**. LDHA mutated lysine (K) to glutamic acid (E) at K118 residue increased LDHA activity. The four lysines at K5, K76, K118, and K232, up-regulated succinylated K residues in PC-3M, were mutated to E to mimic the negatively charged succinyl-lysine modification. After PC-3 cells were transfected with LDHA-mutant using Lipofectamine 3000, mutated LDHA activities were determined. **F**. The mutation K to E of LDHA at K118 promotes cell migration and invasion. Cell migration and invasion of PC-3 cells expressed LDHA-K118E were analyzed by wound-healing and Transwell invasion assays. Representative images of invaded cells (200×). Cells that invaded through Matrigel were stained with crystal violet. Bar, 25 μm. Data are mean ± SE. *p < 0.05, **p < 0.01 vs. mock; #p < 0.05, ##p < 0.01 vs. LDHA-WT. See also **Figures S5** and **S6**.

Next, a study was conducted to identify the lysine location that is the exact succinylation that regulates LDH activity. Seven succinylated sites in LDHA were identified through global succinylome studies in PC-3 and PC-3M cells at K5, K76, K81, K118, K126, K232, and K243 (**Figure 3D**). The locations of known acetylation and ubiquitylation in LDHA were marked with succinylation [34, 35]. In PC-3M cells, seven succinylated sites of LDHA were detected of which four sites including K5, K76, K118, and K232 increased more than twice compared to PC-3 cells. Each *mz/mz* spectrum was verified manually (**Figure S6**). Succinylation at K5 and K118 in LDHA were known sites in a previous study; however, no quantitative studies or functional studies of succinylation were conducted in disease situations [11]. Therefore, the increased four Ksu sites of LDHA in PC-3M cells could be suggested as a substrate for SIRT5.

To find sites that regulate LDH activity, each of the four succinylated sites was mutated individually to glutamic acid (E), representing the same physiological properties in protein succinylation. Point mutations of K5, K76, K118, and K232 to glutamic acid resulted in a significant induction in LDHA succinylation (**Figure 3E**). Among the four mutations, K118E mutated LDHA showed the highest activity that LDH activity can be increased by succinylation of LDHA K118. It was also confirmed that the invasion of K118E mutated LDHA overexpressed cells was increased (**Figure 3F)**. In order to verify succinylation at lysine 118 of LDHA, succinyl-LDHA K118 antibody was newly generated by immunizing rabbits with succinyl-K118 peptide conjugated with a carrier protein. The selectivity of the generated antibody was confirmed by dot-blot assay and competitive immunoblot using synthesized LDHA succinyl-K118 peptide (**Figure S7**).

### LDHA-K118su is the key regulator in advanced PCa

As in previous studies, LDHA activity is regulated by protein modification, such as acetylation at K5 or phosphorylation at Y10 [22, 36]. K5 acetylation of LDHA is known to decrease LDHA level by inducing lysoamal degradation of LDHA in human pancreatic cancer [22]. In the regulation of LDHA lever in cancer, acetylation and succinylation of LDHA may play opposing roles. To study the correlation between acetylation and succinylation in LDHA, PC-3 cells were treated with acetate and succinate, respectively. When acetate was treated to increase LDHA acetylation, LDHA was reduced as in the previous study (**Figure 4A**). In PC-3 cells, succinate treatment increased migration and proliferation as much as PC-3M levels, whereas acetate treatment did not increase migration and proliferation (**Figure S5**). When succinate was treated in the same cell, succinylation was increased at the K118 residue (**Figure 4A**). Succinylation in LDHA-K118 decreased as SIRT5 expression increased in seven types of PCa-related cell lines (**Figure 4B**).

**Figure 4.**
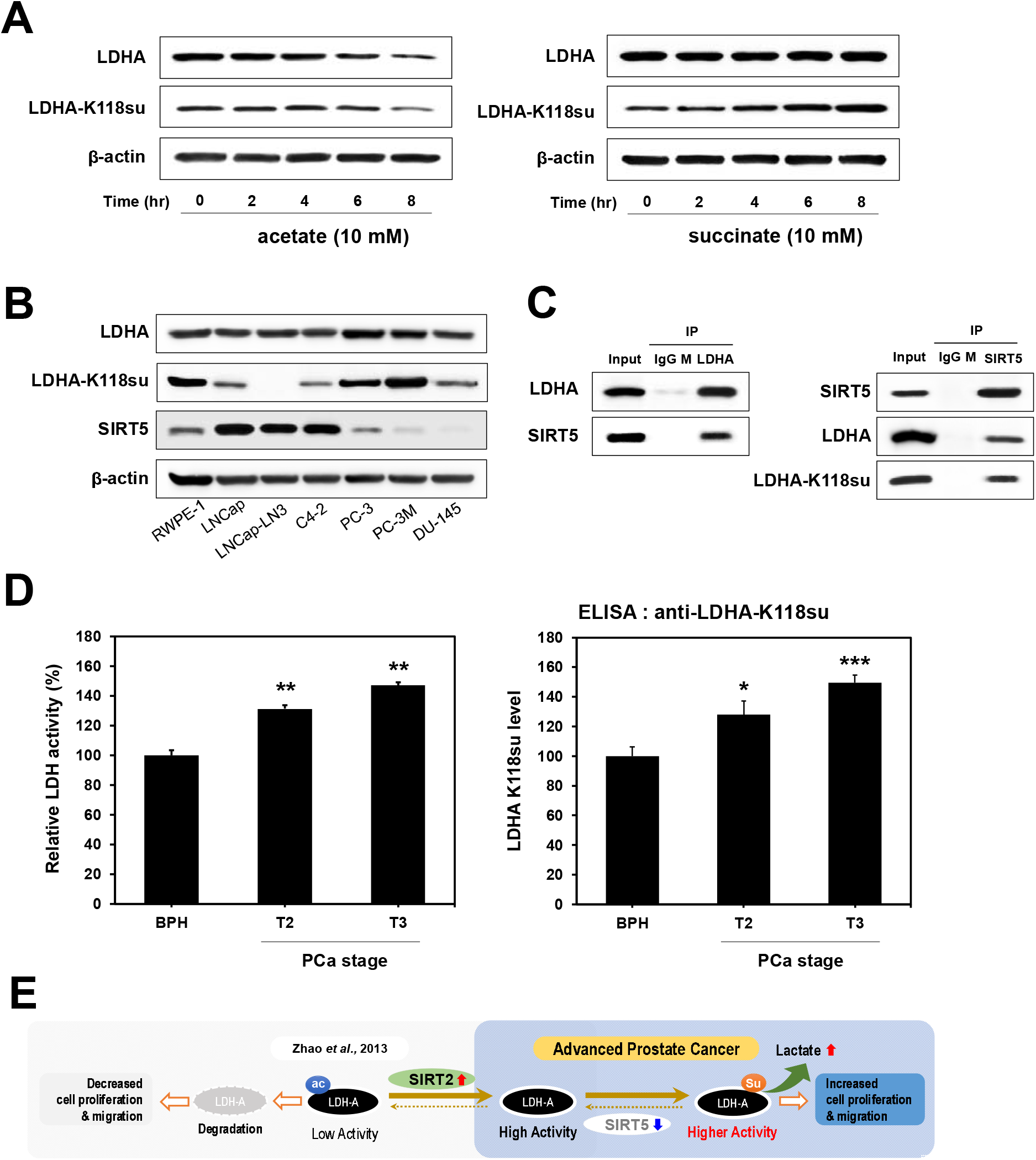
LDHA succinylation promotes PCa progression. **A**. LDHA acetylation induces LDHA degradation. Succinate increased succinylation at the K118 residue of LDHA. The levels of LDHA and LDHA-K118su were determined by Western blot after treatment with acetate or succinate in PC-3 cells for 8 h. **B**. Expression of LDHA, LDHA-K118su, and SIRT5 in seven types of PCa cell lines was evaluated by immunoblot. **C**. SIRT5 interacts with LDHA. The immunoblots showed the IP of SIRT5 with anti-LDHA antibodies in PC-3 cells and IP of LDHA with anti-SIRT5 antibodies in PC-3 cells, respectively. **D**. As PCa progresses, LDHA becomes the K118su residue, and activity increases. LDHA activity was determined in PCa tissues classified by pathological grade (BPH, T2, and T3 grades). The K118su level of LDHA residue in PCa tissue was measured by ELISA using the LDHA-K118su antibody. **E**. Working model of LDHA succinylation at K118 in aggressive PCa cells. LDHA degradation is regulated according to acetylation at the K5 position by SIRT2, according to Zhao et al. 2013. In PCa, the SIRT5 protein level is reduced; the K118 of LDHA as the substrate of SIRT5 is not desuccinylation. Succinylated LDHA is more active than LDHA, so it increases lactate production, leading to PCa progression. Data are mean ± SE. *p < 0.05, **p < 0.01, ***p < 0.001 vs. BPH.

To verify the possibility of interaction between SIRT5 and LDHA, co-IP experiments were performed in PC-3 cells using anti-SIRT5 and anti-LDHA, respectively. In the co-IP experiment, when LDHA and SIRT5 antibodies were used, co-precipitation of SIRT5 and LDHA was demonstrated using immunoblot, respectively (**Figure 4C**). Conversely, when SIRT5 was immunoprecipitated, LDHA was precipitated together. From the above results, it could be estimated that LDHA-K118su is desuccinylated using SIRT5. In patient tissues, as PCa progressed, LDH activity increased significantly compared to BPH in the T2 and T3 grades, which was associated with SIRT5 reduction, as suggested in **Figure 1E** (**Figure 4D**). In **Figure 1E**, total Ksu increased with PCa progression. For more accurate results, when tested with the LDHA-K118su-specific antibody that we produced, LDHA-K118su expression in PCa tissues was also significantly increased according to the stage of PCa progression. In advanced PCa, SIRT5 was decreased and succinylated LDHA increased which was more active than intact LDHA. It has been suggested that succinylated LDHA promotes the progression of prostate cancer (**Figure 4E**).

## Discussion

SIRT5 is involved in cell metabolism, including glycolysis, tricarboxylic acid cycle, fatty acid oxidation, nitrogen metabolism, pentose phosphate pathway, antioxidant defense, and apoptosis [37]. Although SIRT5 has important regulatory roles in tumor progression in the liver and gastric cancer, the role of SIRT5 in cancer has not been extensively studied compared to other SIRTs [38, 39]. Recent studies have shown that SIRT5 is associated with cancer progression through the regulation of succinylation of specific proteins involved in cancer progression [21, 40, 41].Here, SIRT5 levels in PC-3M were significantly lower than in PC-3 cells as well as other androgen-dependent PCa cell lines (LNCaP and LNCaP-LN3). SIRT5 expression and overall survival ratio showed a clear correlation, although not statistically significant. SIRT5 expression was significantly decreased as PCa progressed to T2 and T3, whereas K118su and LDH activity increased. These results led to the speculation that the increase in succinylation of LDHA by the decrease of SIRT5 regulates the aggression of PCa by the increase in LDH activity.

To investigate the molecular mechanism of SIRT5 decrease in PC-3M cells, the expression level of *Sirt5* mRNA was measured in PC-3 and PC-3M cells, however there was no change (**Figure S1C**). Furthermore, a protein synthesis inhibitor (cycloheximide) was treated, and *Sirt5* expression was not affected by cycloheximide (data not shown). In additional experiments, treatment with a proteasomal inhibitor (MG132) or a lysosomal inhibitor (chloroquine) did not alter SIRT5 expression (data not shown). These results indicate that the decrease of SIRT5 in PC-3M cells was not regulated by transcriptional regulation or protein stability. Recently, SIRT5 was reported to be a downstream target of several microRNAs, including miR-299-3p, miR-19b, and miR-3677-3p, which regulate the proliferation, migration, and invasion of cancer cells [42-44]. However, further studies are required to understand and determine the microRNAs related to SIRT5 decrease in PC-3M cells.

LDHA showed a high expression profile and activated status in many cancers [32]. A number of previous studies showed the correlation with LDH activity and tumor growth and metastasis[33, 45-48]. LDHA plays an important role in metastasis and hepatocellular carcinoma cell growth [45]. LDHA was up-regulated in breast cancer and correlated with poor clinical outcomes of breast cancer [46]. When LDHA expression was reduced by small interfering RNA, the progression of lymphoma growth was inhibited [47]. Especially in PCa, high LDHA expression in PC-3 cells induced a favorable tumor microenvironment for tumor progression [48]. As a result, there is a clear correlation that increasing LDH activity increases tumor progression.

The correlation between increased LDHA expression and cancer progression means that LDH activity is important for regulating tumor therapies. However, there are no suitable LDHA inhibitors to use for tumor therapies in the clinic [33]. The regulation of LDH activity in cancer cells is very important for cancer progression, and protein modification plays a key role in regulating LDH activity. As mentioned previously, LDH activity is regulated by protein modification, such as acetylation at K5 or phosphorylation at Y10 [22, 36]. Acetylation at K5 of LDHA in human pancreatic cancer increases the lysosomal degradation of LDHA, and LDH activity was inhibited [22]. In breast and colorectal cancer stem cells, LDHA phosphorylation at Y10 was positively correlated with the progression of metastasis and cell invasion [36, 49].

In this study, we reported a new mechanism in regulating LDHA activity in aggressive PCa, LDHA increases its activity by K118su, and SIRT5 regulates desuccinylation. Overall, SIRT5 increased protein stability due to succinylation and delayed degradation due to succinylation, resulting in increased activity. The mechanism is explained by increasing the lactate of cancer cells according to the increased LDHA activity to promote cancer cell growth [50-52]. Even in PCa cells, decreasing the lactate concentration significantly decreased the growth of cancer cells [52]. Regarding LDHA activity, K118su of LDHA after a decrease in SIRT5 may be related to ubiquitination modified at the same site (**Figure 3D**) [34]. It can be estimated that LDHA activity is maintained by inhibiting degradation because K118su does not proceed with ubiquitylation. Since succinylated LDHA shows higher activity than intact LDHA, a lower SIRT5 indicates an increase in the highly active form of LDHA (**Figure 4E**). A previous study reports no change in the activity of LDHA in SIRT5 KO cell line of colorectal cancer [41]. Although the level of succinylated-LDHA was not evaluated in that study, it was assumed that the process of LDHA succinylation is also a key point like desuccinylation. Since the succinylation of LDHA did not increase in colorectal cancer, it can be assumed that the expression of SIRT5 did not affect the change in LDHA activity. Both desuccinylation by SIRT5 and succinylation by succinyl-transferease should be considered for the potential role of LDHA succinylation as a key regulator in promoting cancer progression.

## Conclusion

PCa is the most common type of genital cancer in men, accounting for the highest number of all new cancer cases, and is the third leading cause of cancer-related deaths in the world. SIRT5 was significantly down-regulated in advanced PCa based on PCa cell lines, and reduced SIRT5 was associated with a decreased survival rate in PCa patients. SIRT5, a representative desuccinylase, regulates the succinylation of proteins in the PCa stage. Through global succinylome analysis, LDHA-K118su regulated by SIRT5 was identified, which increased LDHA activity. As the substrate site of SIRT5, LDHA-K118su significantly increased as PCa progressed and showed a positive correlation with increased migration and invasion of PCa cells. Therefore, LDHA activity, which increased PCa progression, is up-regulated by an increased succinylation at the K118 position and was a substrate for SIRT5. Allowing a new mechanism to inhibit PCa progression was suggested. In further studies, it is necessary to evaluate whether succinylation of LDHA at K118 is a target for the development of novel antitumor drugs.

## Materials and Methods

### Clinical samples of prostate tissues

The biospecimens used in this study were provided by the National Biobank of Korea-Kyungpook National University Hospital (KNUH), a member of the Korea Biobank Network. Data were obtained (with informed consent) under institutional review board-approved protocols. Normal tissues (BPH) and PCa tissues (separated into T2 and T3) were stored at -80°C before use.

### PCa cell line culture

PCa cell lines, including LNCaP, LNCaP-LN3, PC-3, and PC-3M, were purchased by the Korea Cell Line Bank (Seoul, Korea). The detailed experimental methods are described in the supplementary document.

### Western blot analysis

PCa cell lines were lysed using radioimmunoprecipitation assay buffer (Thermo, Waltham, MA, USA). Protein concentrations were measured by BCA assay (Thermo). Samples were separated via 10% sodium dodecyl sulfate (SDS)-polyacrylamide gel electrophoresis for 90 min and transferred to polyvinylidene fluoride blotting membranes (0.2 µm pore size; GE Healthcare, Chalfont St. Giles, UK) for 2 h. The membrane was blocked in 5% bovine serum albumin for 2 h. Primary antibodies against SIRT1-SIRT7 (Abcam, Cambridge, UK) and succinyl-lysine (PTM Biolabs, Chicago, IL, USA) were used to capture SIRT1-SIRT7 proteins overnight at 4°C. After incubation with horseradish peroxidase (HRP)-conjugated secondary antibody for 1 h, the membranes were washed in Tris-buffered saline-0.1% Tween 20 (TBS-T) three times for 15 min. ECL Prime Western blotting detection reagent (GE Healthcare) was used to visualize the blots.

### Wound-healing and Transwell invasion assays

A wound-healing assay was performed to observe the migratory capacity of PC-3 and SIRT5-KO cells. Cells (0.4 × 10^6^ per well) were seeded in each well of a 6-well plate (Corning, Corning, NY, USA) and cultured to ∼90% confluence. A sterilized 1000 μL tip was used to scratch the cell monolayer artificially, and debris was then removed by washing with phosphate-buffered saline (PBS). Next, wounded cells were observed under a microscope at 12 h intervals and imaged with a Leica Las EZ (Olympus, Tokyo, Japan).

To evaluate the invasiveness of PC-3 and PC-3 SIRT5-KO cells, a Transwell invasion assay was performed with 200 μg/mL Matrigel matrix (Corning) and 8 μm pore-permeable supports (Corning). Cell samples were resuspended in serum-free medium and seeded (5 × 10^4^) in each upper chamber, and the medium containing 10% fetal bovine serum was added into the lower chamber. After 24 h, a cotton swab was used to remove the non-invading cells on the upper chamber. Cells that migrated through the Matrigel matrix were fixed with 4% paraformaldehyde for 10 min and stained with 0.5% crystal violet (Sigma-Aldrich, St. Louis, MO, USA) for microscopy. Finally, cells attached to the membrane were added. Optical density was measured at 570 nm.

### Construction of SIRT5-KO cell lines using the CRISPR/Cas9 gene editing system

We used the CRISPR/Cas9 system to knock out SIRT5 from cell lines. PC-3 cells were transfected with either the control or *Sirt5* CRISPR/Cas9 plasmids according to the manufacturer’s instructions (Sigma-Aldrich). The transfection efficiency was determined by a green fluorescent protein (GFP) signal. After 24 h, GFP-expressing PC-3 cells were sorted by FACS Aria III (BD Life Sciences, Franklin Lakes, NJ, USA) and maintained with complete medium. Sirt5 deletion by the CRISPR/Cas9 *Sirt5* but not the control plasmid was confirmed by Western blot.

### Proliferation assay

Cells (1.5 × 10^4^ per well) were cultured in a 96-well plate in complete RPMI medium. After 24 h, proliferation was determined by CCK-8 (Dojindo, Tokyo, Japan). The CCK-8 solution was added to the plate and incubated for 30 min in a humidified incubator. The plate was measured at 420 nm after the appropriate color change was observed.

### Global IP for comparative succinylome analysis to enrich K118su

Anti-succinyl-lysine antibody-conjugated agarose beads (40 μL) were prewashed with 1 mL PBS followed by centrifugation at 1000×*g* for 1 min at 4°C. To enrich lysine succinylated (K118su) peptides, 5 mg desalted peptides were dissolved in 1 mL NETN buffer [50 mM Tris-HCl (pH 8.0), 100 mM NaCl, 1 mM EDTA, 1% NP-40] and incubated on a rotator at 4°C for 16 h. The beads were washed three times with 1 mL NETN and twice with 1 mL ETN [50 mM Tris-HCl (pH 8.0), 100 mM NaCl, 1 mM EDTA] and 1 mL liquid chromatography/mass spectrometry (LC/MS)-grade water using centrifugation at 1000×*g* for 1 min at 4°C in between washes. The enriched K118su peptides were eluted in 70 μL of 0.15% trifluoroacetic acid (TFA) by gentle mixing. After centrifugation at 1000×*g* for 1 min, 50 μL of 0.15% TFA were added to the beads to improve the recovery efficiency. Finally, 120 μL of the enriched K118su peptides were dried in a speed-vacuum system and then desalted twice using a C18 zip tip. Samples were analyzed using LTQ-velos Orbitrap connected to the Eksigent nanoLC system at Mass Spectrometry Convergence Research Center, and detailed analysis methods are described in the supplement information.

### Site-directed mutagenesis

Human LDHA plasmid was purchased from Origene (MD, USA). Succinylation mutants of LDHA were generated using site-directed mutagenesis kit (GeneAll, Seoul, Korea) by converting each lysine residue (K5, K76, K118, and K232) to glutamic acid (codon change from AAG or AAA to GAG or GAA). Mutation was confirmed by DNA sequencing conducted at Cosmogenetech (Seoul, Korea).

### Transient transfection and in vitro migration assay

To express exogenous proteins, PC-3 cells were mock-transfected or transfected with 0.5 to 4 μg LDHA-WT or LDHA-mutant using Lipofectamine 3000 (Invitrogen, Thermo Fisher Scientific, Bremen, Germany). For the migration assay, cell migration was analyzed using Transwell migration assay chambers (BD Life Sciences). PC-3 cells were transfected with 4 μg plasmid DNA and incubated for 24 h. Then, 1.5 × 10^5^ cells were transferred to Transwell to perform migration assay and incubated for 24 h on Transwell.

### LDH activity colorimetric assay

LDH activity was performed as described (K726-500, BioVision, Milpitas, CA, USA). In brief, 0.1 g tissues or 1×10^6^ cells were homogenized in cold assay buffer and then centrifuged at 10,000×*g* for 15 min at 4°C. The supernatants were used assay. Fifty microliters of the sample were added to a 96-well plate. Then, 50 μL of the reaction mix were added. OD_450 nm_ was measured at time 0 and then measured again after incubation at 37°C for 30 min and then calculated according to the kit protocol.

### Development and characterization of K118-specific antibodies

The polyclonal antibody specific for succinylated LDHA at K118 (anti-succinylated K118-LDHA) was produced in collaboration with PTM Biolabs. Rabbits were immunized with succinylated LDHA peptides (CRNVNIF-Ksu-FIIPNVVK and RKRNVNIF-Ksu-FIIPNC). Antisera from immunized rabbits were first depleted with the unmodified peptide and then affinity purified using the succinylated K118 peptides.

### Data and statistics

All experiments were repeated at least three times. Data were expressed as mean ± standard error (SE). To determine the significant differences, statistical analysis was performed using IBM SPSS Statistics version 21. p < 0.05 was considered statistically significant.

## Authors’ contributions

OKK, TGK, BHP and SL conceived and digesned the experiments. OKK, IHB, SYC, JMJ, AN, YG and SSC performed the experiments. OKK, SHK and SL analyzed the date. YC, JNL, YH, EJB, TGK and BHP contributed reagents/materials/analysis tools. OKK, SYC and SL wrote the paper. TGK, BHP and SL revised the manuscript. All authors read and approved the final manuscript.

## Acknowledgements

This work was supported by the National Research Foundation of Korea (NRF) and funded by the Korean Government (MSIP; grant Nos. NRF-2018R1D1A1A02043591, 2019R1H1A1079839, 2020M3A9B6037812, and 2020R1A2B5B03002344). It was also supported by KBRI (21-BR-02-04).

## Supplementary material

**Figure S1. Reduction of SIRT5 in PCa cells A**. Immunostaining of SIRT5 indicates decreased SIRT5 in PC-3M and PC-3 SIRT5-KO compared to PC-3 cells by confocal microscopy. **B**. The level of SIRT5 level was decreased in the cytoplasm and mitochondria in PC-3M and PC-3 SIRT5-KO cells. Fractionation controls were HSP90 (cytosol), CALR (mitochondria), and histone H3 (nucleus). **C**. The level of *SIRT5* mRNA is the same in PC-3 and PC-3M cells. The *SIRT5* level was determined by RT-qPCR in two cell lines. **D**. SIRT5 and *SIRT5* knockdown inhibitor promotes cell migration. Representative images from the wound-healing assay of migrated cells at 0 and 48 h after treatment with SIRT5 selective inhibitor and PC-3, PC-3 SIRT5-KO, and PC-3M cells. **E**. *SIRT5* knockdown cells are depleted only at the SIRT5 level. The levels of seven SIRT5 were analyzed by Western blot in PCa cells (Normal, non-treated PC-3 cells; MOCK, Cas9-vehicle-treated PC-3 cells; PC-3 SIRT5-KO, PC-3 cells knocked down for SIRT5). **F**. *SIRT5* knockdown cells promote cell proliferation. The proliferation of normal, control, PC-3 SIRT5-KO, and PC-3M cells was analyzed by CCK-8. **G**. The level of four lysine acylation was not significantly different in PCa cell lines.

**Table S1. Relationship between clinical characteristics and SIRT5 in PCa patients.** The SIRT5 expression level in clinical cancer tissue showed the most significant correlation with T stage than other clinical indicators.

**Figure S2. Overall survival in PCa patients by SIRT5 level in PCa tissue**

**Figure S3. Global succinylome in PC-3M cells A**. The Pearson correlation coefficients of the ratio of proteins and Ksu peptides are 0.89 and 0.71, respectively. **B**. Characterization of lysine succinylation in prostate cancer using functional enrichment of gene ontology (GO) analysis. **C**. Twelve succinylated proteins overlapped between extracellular and cytoplasm terms. **D**. Succinylation was found in seven lysine residues of LDHA, and the level of four of which increased in PC-3M cells.

**Table S2. Identified protein and succinylated peptide list**

**Table S3. Number of lysine succinylation sites and proteins in PCa cells**

**Table S4. List of up-regulated Ksu proteins, associated with the extracellular and cytoplasm terms, in PCa cells**

**Figure S4. Immunoblots of LDHA and its succinylation. A**. The LDHA level is the same in PC-3 cell lines. LDHA expression was analyzed by Western blot with an anti-LDHA antibody. **B**. SIRT5 knockdown increases the LDHA succinylation level. The acetyl-LDHA level in PC-3 WT cells was higher than PC-3 SIRT5-KO cells. The level of immunoprecipitated LDHA was measured by direct Western blot using pan-succinyl-lysine and acetyl-lysine antibodies. **C**. The LDHA succinylation level is increased by SIRT5 inhibition. After the treatment with SIRT5 inhibitor, LDHA was immunoprecipitated, and LDHA succinylation was analyzed by Western blot using a pan-succinyl-lysine antibody. **D**. Succinate stimulates the LDHA succinylation level. After treatment with succinate, LDHA was immunoprecipitated, and LDHA succinylation was analyzed by Western blot using a pan-succinyl-lysine antibody.

**Figure S5. Increased the migration and proliferation of PC-3 in hypersuccinylation. A**. Succinate increased the cell migration. Representative images from the wound-healing assay of migrated cells at 0 and 48 h after treatment with succinate and acetate to PC-3 cells. **B**. Succinate increased the cell proliferation. The proliferation of PC-3 after succinate and acetate treatment, and PC-3M cells was analyzed by CCK-8 assay.

**Figure S6. The mz/mz spectra show Ksu at K5, K76, K118, and K232 in LDHA**

**Figure S7. Validation of selectivity of anti-LDHA-K118su A**. Dot-blot assay showed the selectivity of anti-LDHA-K118su. Consistent with the ELISA result, the purified antibody showed very good immune-response with the modified peptides. The detection limit is lower than 4 ng. The signal doesn’t recognize the control peptide. **B**. Competitive assay of anti-LDHA-K118su in PC-3 and PC-3M cell lines using synthetic LDHA, succinyl-K118 peptides.

