## Supplemental information for "SIRT5 is the desuccinylase of LDHA as novel cancer metastatic stimulator in aggressive prostate cancer"

^#^ Equal contribution.

^*^ Corresponding author.

**Supporting Methods**

**Materials.** For screening of lysine acylation in PCa, primary antibodies specific to pan anti-succinyl-lysine antibody (PTM-401; PTM Biolabs), pan anti-acetyl-lysine antibody (PTM-101; PTM Biolabs), pan anti-malonyl-lysine antibody (PTM-901; PTM Biolabs), pan anti-glutaryl-lysine antibody (PTM-1151; PTM Biolabs), and pan anti-3-hydroxy-butyryl-lysine antibody (PTM-1201; PTM Biolabs) were purchased. HRP-linked mouse and rabbit secondary antibodies were obtained from Cell Signaling Technology (Beverly, MA, USA).

**PCa cell-line culture.** The PCa cell lines, including LNCaP, LNCaP-LN3, PC-3, and PC-3M, were purchased by the KCLB. RWPE1 as a normal prostate cell line was obtained from the American Type Culture Collection (Manassas, VA, USA). Cancer cell lines were cultured in RPMI 1640 medium supplemented with 10% FBS and 1% penicillin G and streptomycin as antibiotics at 37°C in an atmosphere of 5% CO_2_ in a humidified incubator. The normal prostate cell line was grown in keratinocyte SFM (Gibco, Thermo Fisher Scientific, Bremen, Germany) with a human keratinocyte growth supplement kit (Gibco, Thermo Fisher Scientific), including human recombinant epidermal growth factor and a bovine pituitary extract.

**Sandwich ELISA.** To measure the level of SIRT5 and succinyl-lysine (Ksu) in prostate tissues, 50 μL/well of mouse monoclonal antibody for SIRT5 and Ksu diluted with 0.2 M sodium bicarbonate (2 μg/mL) was used to coat a sterilized 96-well polystyrene plate (Corning) for 2 h. Next, the plate was washed three times with TBS-T and incubated with 3% BSA in TBS-T overnight at 4°C. Normal prostate tissues and PCa tissues lysed with RIPA buffer were incubated for 5 h at room temperature (RT) and then washed again. Then, 50 μL/well detection antibody (SIRT5 rabbit polyclonal antibody diluted with 3% BSA) was added for overnight incubation at 4°C. After washing the plate three times, a secondary antibody (HRP-conjugated rabbit antibody) was added for 2 h, followed by five washes. Finally, the TMB substrate (Thermo Fisher Scientific, Waltham, MA, USA) was added to each well to visualize the appropriate color. A stop solution (Thermo Fisher Scientific) was used to halt the color change to ensure precise measurement. The plate was measured for optical density at 450 nm.

**Immunofluorescence staining.** The cells were fixed with 3.2% paraformaldehyde (Electron Microscopy Sciences, Hatfield, PA, USA) for 30 min at RT. The cells were permeabilized with 0.1% Triton X-100 in PBS (PBST) and then blocked with 10% normal goat serum (Vector Laboratories, Burlingame, CA, USA) in PBST for 1 h at 37^o^C. The cells were incubated with primary antibodies against SIRT5 (Abcam) and COX IV(Abcam) diluted in PBST containing 2% normal goat serum at 4^o^C overnight and then washed thrice for 10 min. Subsequently, the cells were then incubated with Alexa 488- or 568-conjugated secondary antibodies for 3 h at RT followed by staining with 4′,6-diamidino 2-phenylindole at RT for 8 min. The samples were mounted on slides using Vectashield mounting medium and imaged using a confocal microscope (Leica Biosystems, Buffalo Grove, IL, USA) using 600× objective for imaging.

**Real-time quantitative polymerase chain reaction (RT-qPCR).** Total RNA was isolated from PC-3 and SIRT5-KO cells using an RNA Cell Miniprep kit (Promega, Fitchburg, WI, USA) according to the manufacturer’s instructions. cDNA synthesis was performed using a GoScript Reverse Transcription kit (Promega) and then used as a qPCR template with FASTStart Essential DNA Green Master mix (Roche, Basel, Switzerland). The primer sequences were as follows: IL-1β forward 5′-CTTCGAGGCACAAGGCACAA-3′ and reverse 5′-TTCACTGGCGAGCTCAGGTA-3′ and ACTB forward 5′-TGCGCCGTTCCGAAAGTT-3′ and reverse 5′-GCGCCGCTGGGTTTTATAG-3′.

**Stable isotope labeling by amino acids in cell culture (SILAC).** For SILAC experiments, PC-3 and PC-3M cells were labeled with light or heavy amino acids in SILAC RPMI 1640 medium over seven growth passages. SILAC RPMI 1640 medium was supplemented with 48 mg/L lysine and 200 mg/L arginine (SILAC “light”) or ^13^C_6_^15^N_2_-lysine and ^13^C_6_^15^N_4_-arginine (SILAC “heavy”). Biological replicates were harvested 2 weeks apart from the identical cell culture. After confirming that the SILAC labeling efficiency was >98%, the cells were lysed with RIPA buffer, and lysates were quantified by BCA assay. We mixed the two cell lines equally (300 μg each) and performed reduction and alkylation using 15 mM DTT and 60 mM IAA. Overnight trypsin digestion at 37°C was used to digest proteins into peptides. Then, high-pH reverse-phase fractionation and off-gel fractionation were performed according to the manufacturer’s protocol.

**Sample preparation for proteomic analysis. Sample preparation and in-solution tryptic digestion. Dishes of SILAC-labeled cancer cells were washed twice with PBS on ice and then scraped into 4% SDS lysis buffer with halt protease inhibitor cocktail, 4 M sodium butyrate, and 2 M nicotinamide as HDAC inhibitors. To ensure complete protein extraction, the cell lysates were disrupted by sonication at 4°C for 1 min and heated for 5 min at 98°C. Any unbroken cells were removed by centrifugation at 16,000×*g* for 10 min at 4°C, and the supernatant was transferred to a low-protein binding E-tube. Proteins from PC-3 and PC-3M cells were measured using a BCA protein assay kit. The same amount of proteins from PC-3 and PC-3M cells were carefully mixed for SILAC-based quantitative proteomics. The combined proteins were sequentially reduced and alkylated with 15 mM DTT at 56°C for 30 min and 60 mM IAA for 30 min at RT in the dark. The detergent and chemical reagents present in the protein mixture were removed after protein precipitation with 10% trichloroacetic acid for 4 h at 4°C. After centrifugation at 12,000×*g* for 10 min at 4°C, the protein pellets were washed twice with 20°C acetone and then resuspended in 50 mM ABC buffer using sonication for 5 min on ice. To generate the peptides, the SILAC proteins were digested with trypsin at 37°C overnight on a rotator. A 10% solution of TFA was added to the peptides to a final concentration of 1% to stop the trypsin reaction. The reaction mixture was centrifuged at 16,000×*g* for 5 min to remove the enzyme and any undigested protein. Before immunoprecipitation to assess lysine succinylation, the peptides were cleaned using a C18 Sep-Pak according to the manufacturer’s instructions. The C18 3cc Sep-Pak was wetted with 3 mL of 100% ACN and 2 mL of 50% ACN. For equilibration, the Sep-Pak was washed three times by adding 2 mL water containing 0.1% TFA. The peptides were loaded slowly into the cartridge and then desalted three times using 2 mL water containing 0.1% TFA. The samples were eluted twice with 1 mL of 75% ACN containing 0.1% TFA each time and for a third time to maximize peptide recovery. The purified samples were dried using a speed-vacuum system. A quantitative colorimetric peptide assay was used to measure the amount of peptide in the samples.**

**LC-MS/MS analysis. Peptide fractionation for in-depth quantitative proteomics. To improve protein identification, the SILAC peptides were separated by two different processes, high-pH reverse-phase fractionation and 3100 OFFGEL fractionation. The fractionation of peptides was performed according to the manufacturer’s procedures. Briefly, peptides (100 μg) were made of 0.1% TEA in ACN and separated using high-pH reverse-phase fractionation kit using eight different elution buffers (5%, 7.5%, 10%, 12.5%, 15%, 17.5%, 20%, and 50% ACN in 0.1% TFA). The eight samples were combined into four samples (e.g., fractions 1 and 5, 2 and 6, etc.). Peptides (200 μg) were separated by OFFGEL fractionation using high-resolution 24-well frame IPG strips (pH 3–10). These were also combined into 12 samples derived from 24 fractions. All fractionated samples were desalted using a C18 zip tip and completely dried using a speed-vacuum system before further analysis.**

**All samples were dissolved in solvent A (98% water in 0.1% FA). In particular, IP samples were centrifuged at 16,000×*g* for 5 min to remove any remaining beads. Samples were analyzed using a high-resolution, accurate MS connected to the Eksigent nanoLC system at Mass Spectrometry Convergence Research Center. Peptide separation was conducted using a home-made C12 reverse-phase analytical column with a linear gradient of 0–23% solvent B (100% ACN in 0.1% FA) for 95 min, 23–90% solvent B for 9 min, and 90% solvent B for 6 min at a sustained flow rate of 300 nL/min. The LTQ-velos Orbitrap was operated in the top 20 data-dependent acquisition (DDA) mode. MS data were collected using the following settings: electrospray source voltage 1.8 kV and a 300°C capillary temperature; FTMS 300–1800 m/z range with 60,000 resolution (*m/z* 400); collision-induced dissociation mode at 28% NCE and an isolation width of 1.7 *m/z*; and 500 minimum signals were required for DDA. Lock mass ion from ambient air (*m/z* 445.120024) was enabled to improve mass accuracy.**

**Bioinformatics. Data analysis and bioinformatics for the succinylome. To identify proteins and Ksucc proteins, MS/MS spectra were processed using MaxQuant 1.5.1.0 to query the UniProtKB human database (including 71,772 protein sequences; 19029910). The search parameters used a full mass error of 20 ppm and an MS/MS error of 0.5 Da. Trypsin/p was used as the digestion enzyme, allowing for two missing cleavages. The fixed modification was carbamidomethylation of cysteine. For the proteomic search, the oxidation of methionine and acetylation of the N-terminus were set as the variable modifications. For succinylation analysis, Ksu was added as the variable modification. For quantification by SILAC, Lys8 (^13^C_6_^15^N_2_) and Arg10 (^13^C_6_^15^N_4_) were specified as the heavy labels (H) and Lys0 (^12^C_6_^14^N_2_) and Arg0 (^12^C_6_^14^N_4_) were set as the light labels (L). The SILAC pairs (H/L) were detected and quantified from full MS using MaxQuant software. The ratio determined the H/L and 2 of the minimum ratio count. The other parameters for MaxQuant were set to the default values.**

**The search results were filtered with a false discovery rate of below 0.01, a MaxQuant score more than 40, discarding potential contaminants, and those only identified by a site modification. Also, Ksu sites were specified using a site localization probability of > 0.75. It should be noted that the Ksucc ratio was normalized to protein levels. All ratios of proteins and lysine succinylated peptides were converted to a log_2_ scale.**

**Protein Co-immunoprecipitation.** LDHA (Cell Signaling Technology), SIRT5 (abcam)or mouse IgG (Santa Cruz Biochemicals, Dallas, TX, USA) antibody and protein G magnetic nanobeads (Bioneer, Daejeon, Korea) were mixed and incubated in a rotator for 30 min at RT and then washed with nanobeads twice. At least 500 ng of protein were mixed with nanobeads and then incubated in a rotator for 1 h at RT and then washed with nanobeads twice. The elution buffer was added and eluted in a vortex for 5 to 10 min and then incubated for 10 min in 4% SDS at 95°C.

**Indirect ELISA for determination of LDHA-K118su in PCa tissues.** Prostate tissue was lysed with RIPA buffer (Thermo), and 1 μg tissue sample was mixed with 50 μL coating buffer (0.2 M sodium bicarbonate, pH 9.4) and incubated for 7 h at RT in a 96-well plate (Corning) and then washed three times with wash buffer (TBS-T pH 7.2) and incubated overnight at 4°C in blocking buffer (3% BSA in TBS-T). Primary antibodies against LDHA-K118su (CTM-212; PTM Biolabs) were diluted 1:1000, added to each well, and incubated at RT for 7 h and then washed three times with wash buffer. HRP-conjugated rabbit antibody was diluted 1:2000, added to each well, and incubated for 3 h and then washed five times with wash buffer. TMB substrate solution (Thermo Fisher Scientific) was added to each well. After 10 to 15 min, when the color was changed properly, a stop solution (Thermo Fisher Scientific) was added, and OD_450 nm_ was measured.

**Supporting Results**

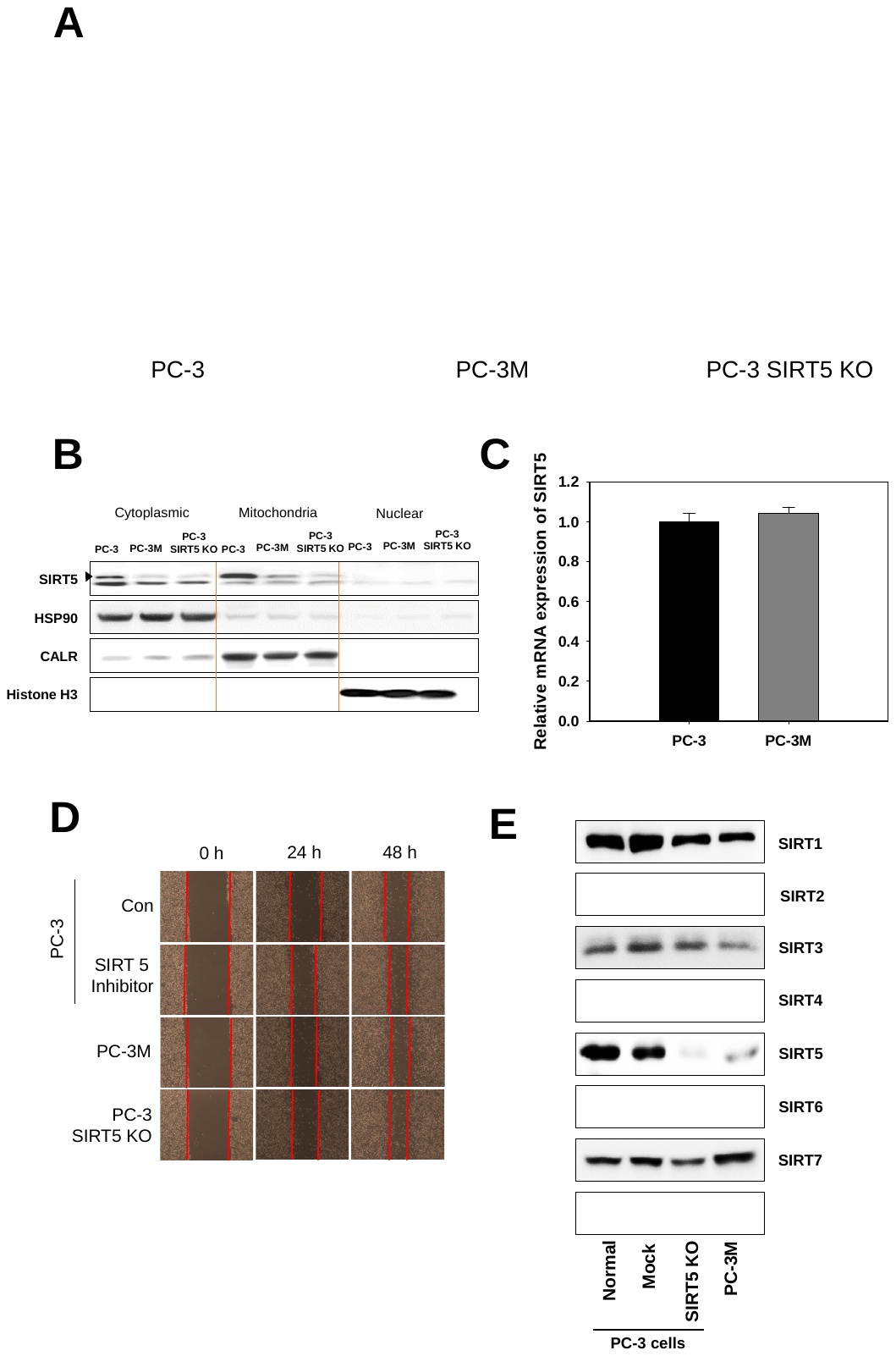

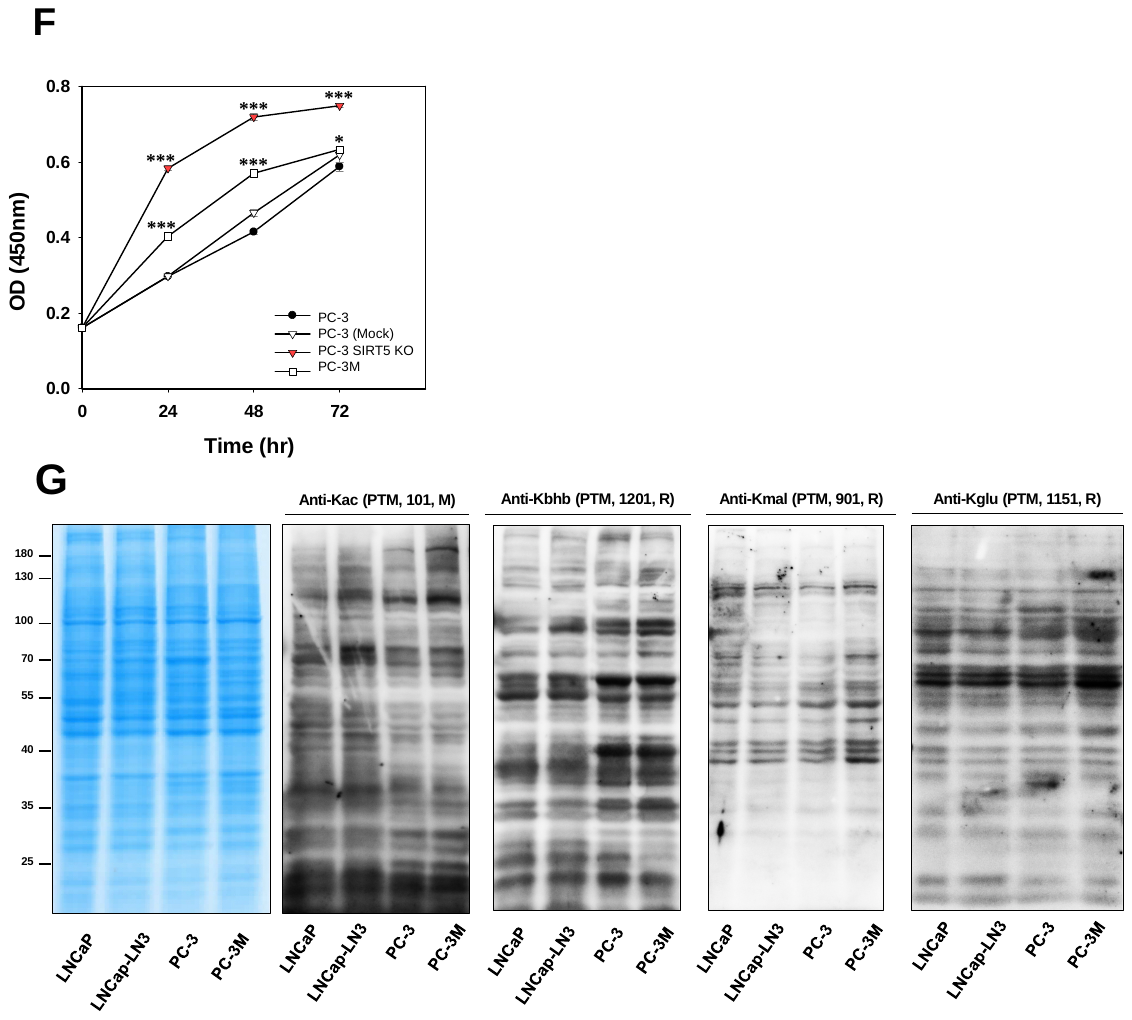

**Figure S1. Reduction of SIRT5 in PCa cells**

**A.** Immunostaining of SIRT5 indicates decreased SIRT5 in PC-3M and PC-3 SIRT5-KO compared to PC-3 cells by confocal microscopy. **B.** The level of SIRT5 level was decreased in the cytoplasm and mitochondria in PC-3M and PC-3 SIRT5-KO cells. Fractionation controls were HSP90 (cytosol), CALR (mitochondria), and histone H3 (nucleus). **C.** The level of *SIRT5* mRNA is the same in PC-3 and PC-3M cells. The *SIRT5* level was determined by RT-qPCR in two cell lines. **D**. SIRT5 and *SIRT5* knockdown inhibitor promotes cell migration. Representative images from the wound-healing assay of migrated cells at 0 and 48 h after treatment with SIRT5 selective inhibitor and PC-3, PC-3 SIRT5-KO, and PC-3M cells. **E**. *SIRT5* knockdown cells are depleted only at the SIRT5 level. The levels of seven SIRT5 were analyzed by Western blot in PCa cells (Normal, non-treated PC-3 cells; MOCK, Cas9-vehicle-treated PC-3 cells; PC-3 SIRT5-KO, PC-3 cells knocked down for SIRT5). **F**. *SIRT5* knockdown cells promote cell proliferation. The proliferation of normal, control, PC-3 SIRT5-KO, and PC-3M cells was analyzed by CCK-8. **G**. The level of four lysine acylation was not significantly different in PCa cell lines.

**Table S1.** **Relationship between clinical characteristics and SIRT5 in PCa patients.** The SIRT5 expression level in clinical cancer tissue showed the most significant correlation with T stage than other clinical indicators.

| Variables | Categories | n | SIRT5 level | | *p* |
| --- | --- | --- | --- | --- | --- |
|  |  |  | High | Low |  |
| Age | <66 | 12 | 7 | 5 | 0.209 |
|  | >65 | 13 | 4 | 9 |  |
| Weight | <69 | 14 | 7 | 7 | 0.674 |
|  | >68 | 11 | 4 | 7 |  |
| BMI | <26 | 15 | 8 | 7 | 0.903 |
|  | >25 | 9 | 5 | 4 |  |
| ECOG performance | Yes | 13 | 7 | 5 | 0.014 |
|  | No | 12 | 4 | 8 |  |
| Drinking | Yes | 9 | 7 | 2 | 0.151 |
|  | No | 15 | 4 | 11 |  |
| Smoking | Yes | 11 | 3 | 8 | 0.01 |
|  | No | 14 | 8 | 6 |  |
| Voiding difficulty | Yes | 13 | 10 | 3 | 0.318 |
|  | No | 11 | 3 | 8 |  |
| **pT** | **T2** | **11** | **7** | **4** | **0.003** |
|  | **T3** | **12** | **4** | **8** |  |
| PSA | <9 | 14 | 9 | 5 | 0.013 |
|  | >9 | 11 | 2 | 9 |  |
| Gleason score | 6–7 | 13 | 6 | 7 | 0.596 |
|  | >8 | 9 | 4 | 5 |  |

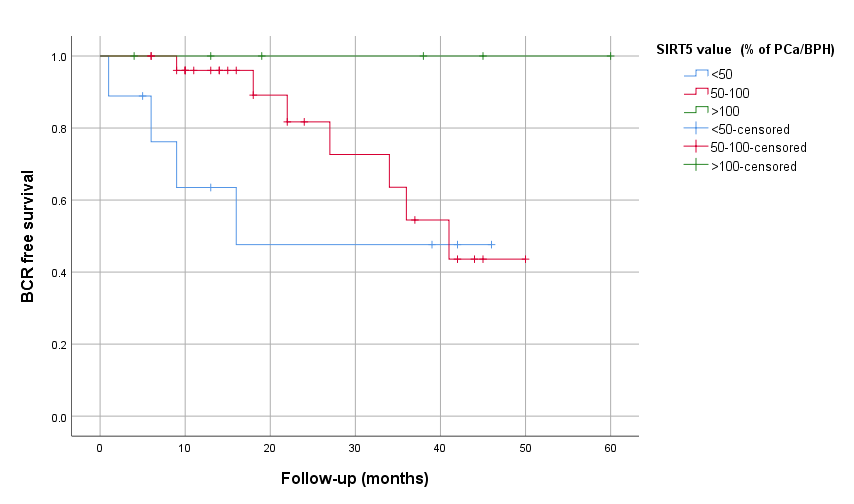

**Figure S2 (related to Figure 1).** Overall survival in PCa patients by SIRT5 level in PCa tissue

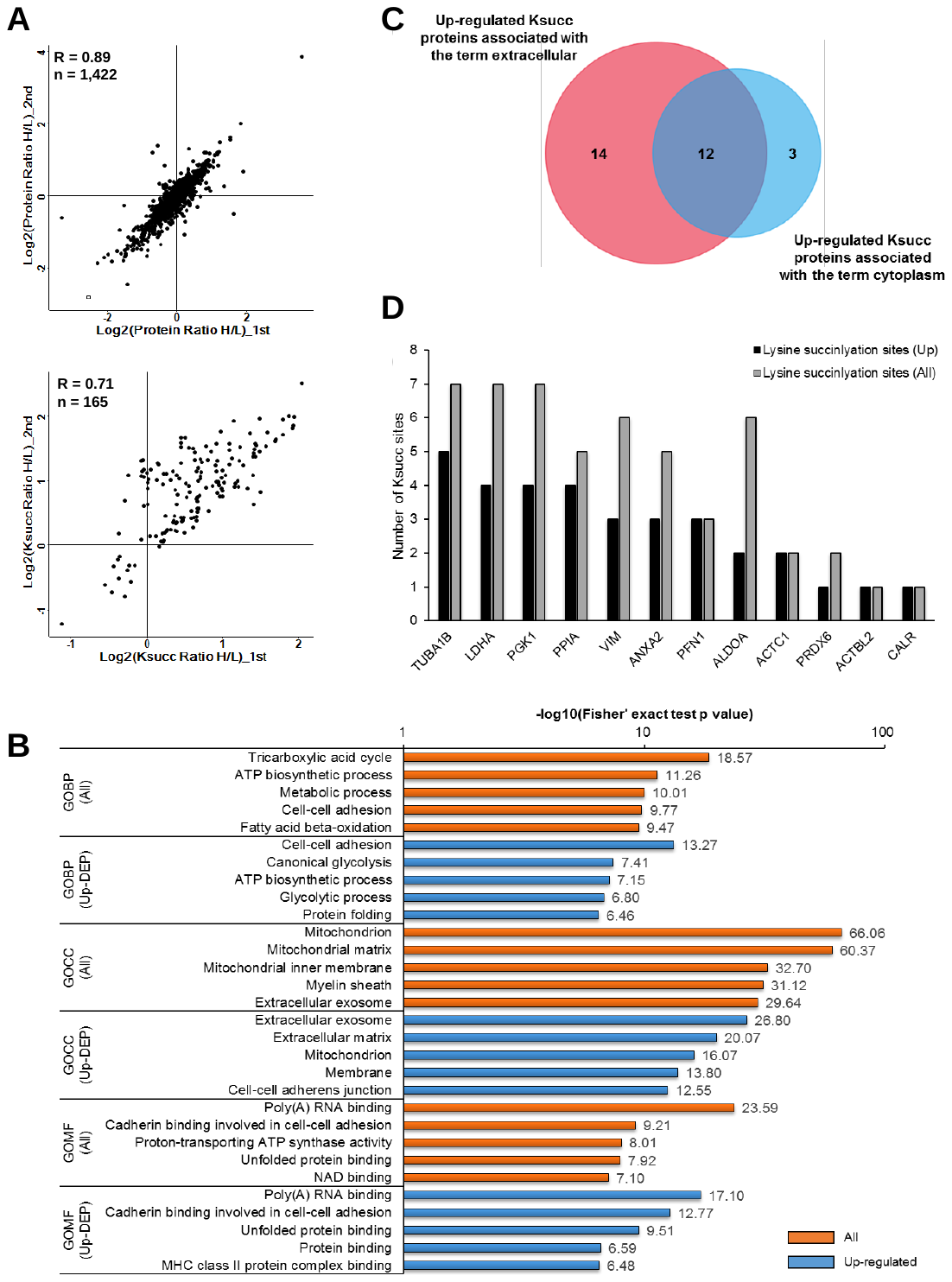

**Figure S3. Global succinylome in PC-3M cells**

**A**. The Pearson correlation coefficients of the ratio of proteins and Ksu peptides are 0.89 and 0.71, respectively. **B**. Characterization of lysine succinylation in prostate cancer using functional enrichment of gene ontology (GO) analysis. **C**. Twelve succinylated proteins overlapped between extracellular and cytoplasm terms. **D**. Succinylation was found in seven lysine residues of LDHA, and the level of four of which increased in PC-3M cells.

**Table S2 (related to Figure 2). Identified protein and succinylated peptide list** (attached separated excel file)

**Table S3. Number of lysine succinylation sites and proteins in PCa cells**

| Succinylome | Identified | Quantified | Normalized quantification | Up-regulated | Down-regulated |
| --- | --- | --- | --- | --- | --- |
| Ksu proteins | 169 | 156 | 136 (80.5%) | 64 (47.1%) | 1 (0.7%) |
| Ksu peptides | 442 | 427 | 403 (91.2%) | 144 (35.7%) | 1 (0.2%) |
| Ksu sites | 448 | 430 | 406 (90.6%) | 144 (35.5%) | 1 (0.2%) |
| Proteomics | 3396 | 2958 (87.1%) | — | 36 (1.2%) | 158 (5.3%) |

**Table S4. List of up-regulated Ksu proteins, associated with the extracellular and cytoplasm terms, in PCa cells**

| Protein ID | Gene name | Score | Modified sequence | Position in peptide | Position in protein | Ksucc ratio | Protein ratio | Normalized Ksucc ratio | Ratio H/L count |
| --- | --- | --- | --- | --- | --- | --- | --- | --- | --- |
| P00338 | LDHA | 217.9 | _(ac)ATLK(su)DQLIYNLLK_ | 4 | 5 | 1.296 | -0.050 | 1.345 | 1 |
| P00338 | LDHA | 72.2 | _TPK(su)IVSGK_ | 3 | 76 | 1.334 | -0.050 | 1.383 | 2 |
| P00338 | LDHA | 152.4 | _NVNIFK(su)FIIPNVVK_ | 6 | 118 | 1.003 | -0.050 | 1.053 | 3 |
| P00338 | LDHA | 63.3 | _EVHK(su)QVVESAYEVIK_ | 4 | 232 | 1.172 | -0.050 | 1.222 | 1 |
| P00558 | PGK1 | 125.7 | _LTLDK(su)LDVK_ | 5 | 11 | 1.177 | 0.163 | 1.014 | 3 |
| P00558 | PGK1 | 79.5 | _IVK(su)DLMSK_ | 3 | 267 | 1.360 | 0.163 | 1.197 | 3 |
| P00558 | PGK1 | 105.6 | _GTK(su)ALMDEVVK_ | 3 | 353 | 1.245 | 0.163 | 1.082 | 1 |
| P00558 | PGK1 | 98.0 | _ALMDEVVK(su)ATSR_ | 8 | 361 | 1.334 | 0.163 | 1.172 | 3 |
| P04075 | ALDOA | 89.1 | _K(su)ELSDIAHR_ | 1 | 14 | 1.879 | 0.521 | 1.358 | 2 |
| P04075 | ALDOA | 120.9 | _VDK(su)GVVPLAGTNGETTTQGLDGLSER_ | 3 | 111 | 1.551 | 0.521 | 1.029 | 2 |
| P07355 | ANXA2 | 165.2 | _TPAQYDASELK(su)ASMK_ | 11 | 115 | 1.526 | 0.473 | 1.053 | 1 |
| P07355 | ANXA2 | 106.4 | _TDLEK(su)DIISDTSGDFRK_ | 5 | 157 | 1.581 | 0.473 | 1.108 | 3 |
| P07355 | ANXA2 | 67.5 | _K(su)LMVALAK_ | 1 | 169 | 1.671 | 0.473 | 1.198 | 1 |
| P07737 | PFN1 | 94.3 | _TK(su)STGGAPTFNVTVTK_ | 2 | 91 | 0.787 | -0.398 | 1.184 | 1 |
| P07737 | PFN1 | 148.5 | _STGGAPTFNVTVTK(su)TDK_ | 14 | 105 | 0.659 | -0.398 | 1.057 | 2 |
| P07737 | PFN1 | 42.8 | _K(su)CYEMASHLR_ | 1 | 127 | 0.697 | -0.398 | 1.095 | 2 |
| P08670 | VIM | 126.4 | _FANYIDK(su)VR_ | 7 | 120 | 0.832 | -0.383 | 1.215 | 3 |
| P08670 | VIM | 127.8 | _K(su)VESLQEEIAFLK_ | 1 | 223 | 0.731 | -0.383 | 1.114 | 1 |
| P08670 | VIM | 124.4 | _TLLIK(su)TVETR_ | 5 | 445 | 1.038 | -0.383 | 1.420 | 3 |
| P27797 | CALR | 60.2 | _NVLINK(su)DIR_ | 6 | 159 | 0.716 | -0.927 | 1.643 | 1 |
| P30041 | PRDX6 | 63.6 | _VVFVFGPDK(su)K_ | 9 | 141 | 1.391 | 0.338 | 1.053 | 1 |
| P62937 | PPIA | 86.4 | _VSFELFADKVPK(su)TAENFR_ | 12 | 31 | 0.670 | -0.392 | 1.061 | 3 |
| P62937 | PPIA | 105.7 | _ALSTGEK(su)GFGYK_ | 7 | 44 | 1.064 | -0.392 | 1.456 | 3 |
| P62937 | PPIA | 48.6 | _TEWLDGK(su)HVVFGK_ | 7 | 125 | 1.041 | -0.392 | 1.433 | 1 |
| P62937 | PPIA | 110.2 | _VK(su)EGMNIVEAMER_ | 2 | 133 | 0.929 | -0.392 | 1.321 | 2 |
| P68032 | ACTC1 | 79.5 | _GILTLK(su)YPIEHGIITNWDDMEK_ | 6 | 70 | 0.992 | -0.211 | 1.203 | 1 |
| P68032 | ACTC1 | 75.8 | _YPIEHGIITNWDDMEK(su)IWHHTFYNELR_ | 16 | 86 | 1.202 | -0.211 | 1.413 | 4 |
| P68363 | TUBA1B | 77.4 | _TIGGGDDSFNTFFSETGAGK(su)HVPR_ | 20 | 60 | 1.169 | -0.163 | 1.333 | 3 |
| P68363 | TUBA1B | 89.5 | _GDVVPK(su)DVNAAIATIK_ | 6 | 326 | 0.941 | -0.163 | 1.104 | 3 |
| P68363 | TUBA1B | 134.4 | _DVNAAIATIK(su)TK_ | 10 | 336 | 0.885 | -0.163 | 1.048 | 3 |
| P68363 | TUBA1B | 121.3 | _LDHK(su)FDLMYAK_ | 4 | 394 | 1.128 | -0.163 | 1.292 | 2 |
| P68363 | TUBA1B | 112.4 | _FDLMYAK(su)R_ | 7 | 401 | 1.143 | -0.163 | 1.306 | 2 |
| Q562R1 | ACTBL2 | 75.6 | _IK(su)IIAPPER_ | 2 | 329 | 0.732 | -0.462 | 1.194 | 4 |

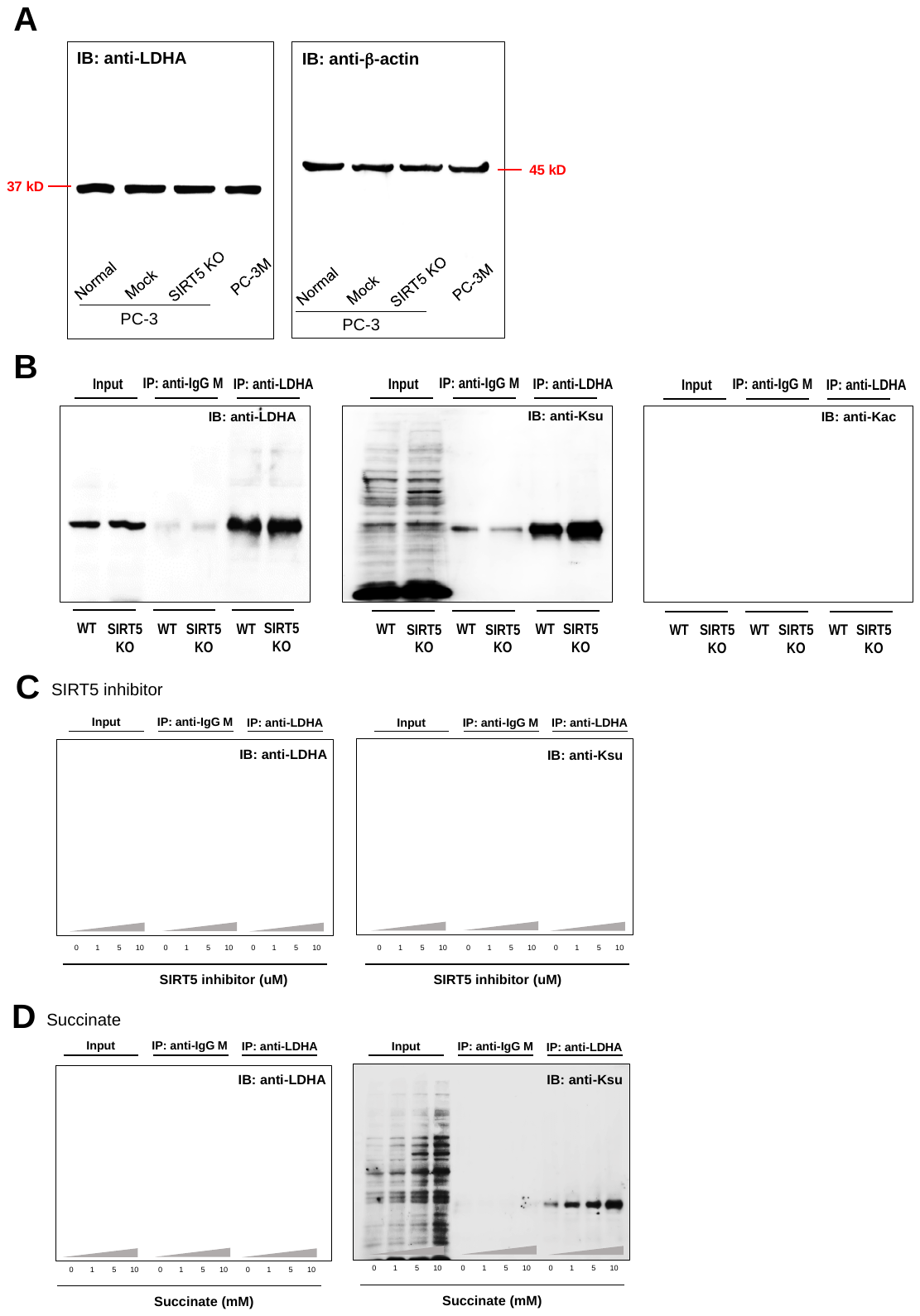

**Figure S4. Immunoblots of LDHA and its succinylation**

**A**. The LDHA level is the same in PC-3 cell lines. LDHA expression was analyzed by Western blot with an anti-LDHA antibody. **B**. SIRT5 knockdown increases the LDHA succinylation level. The acetyl-LDHA level in PC-3 WT cells was higher than PC-3 SIRT5-KO cells. The level of immunoprecipitated LDHA was measured by direct Western blot using pan-succinyl-lysine and acetyl-lysine antibodies. **C**. The LDHA succinylation level is increased by SIRT5 inhibition. After the treatment with SIRT5 inhibitor, LDHA was immunoprecipitated, and LDHA succinylation was analyzed by Western blot using a pan-succinyl-lysine antibody. **D**. Succinate stimulates the LDHA succinylation level. After treatment with succinate, LDHA was immunoprecipitated, and LDHA succinylation was analyzed by Western blot using a pan-succinyl-lysine antibody.

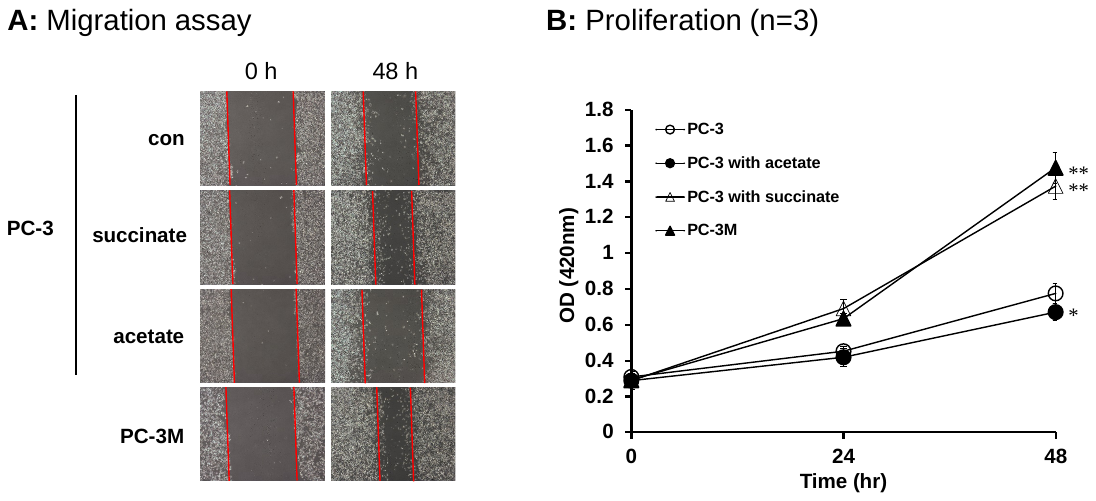

**Figure S5. Increased the migration and proliferation of PC-3 in hypersuccinylation**

**A**. Succinate increased the cell migration. Representative images from the wound-healing assay of migrated cells at 0 and 48 h after treatment with succinate and acetate to PC-3 cells. **B**. Succinate increased the cell proliferation. The proliferation of PC-3 after succinate and acetate treatment, and PC-3M cells was analyzed by CCK-8 assay.

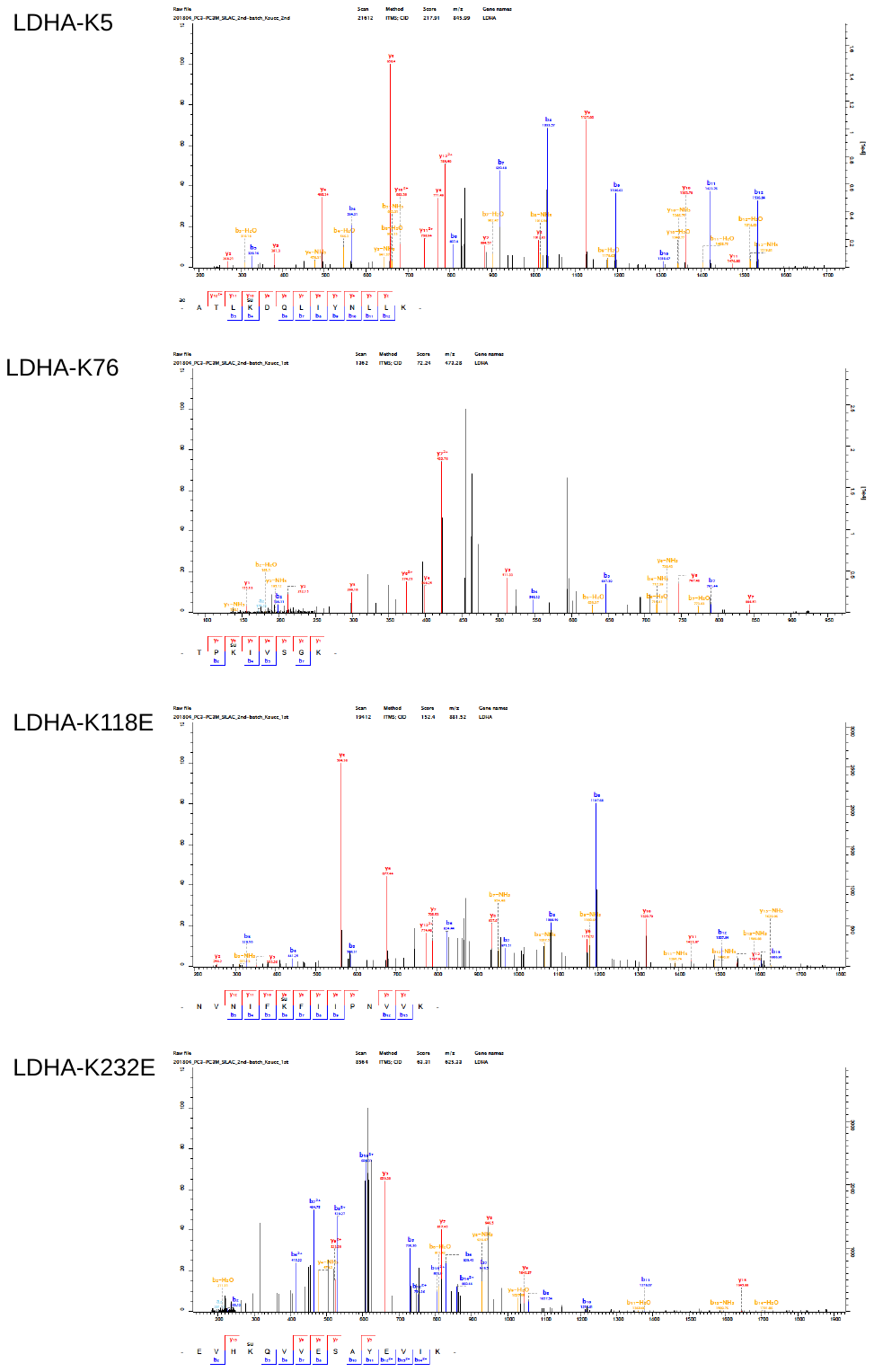

**Figure S6 (related to Figure 3). The *mz/mz* spectra show Ksu at K5, K76, K118, and K232 in LDHA**

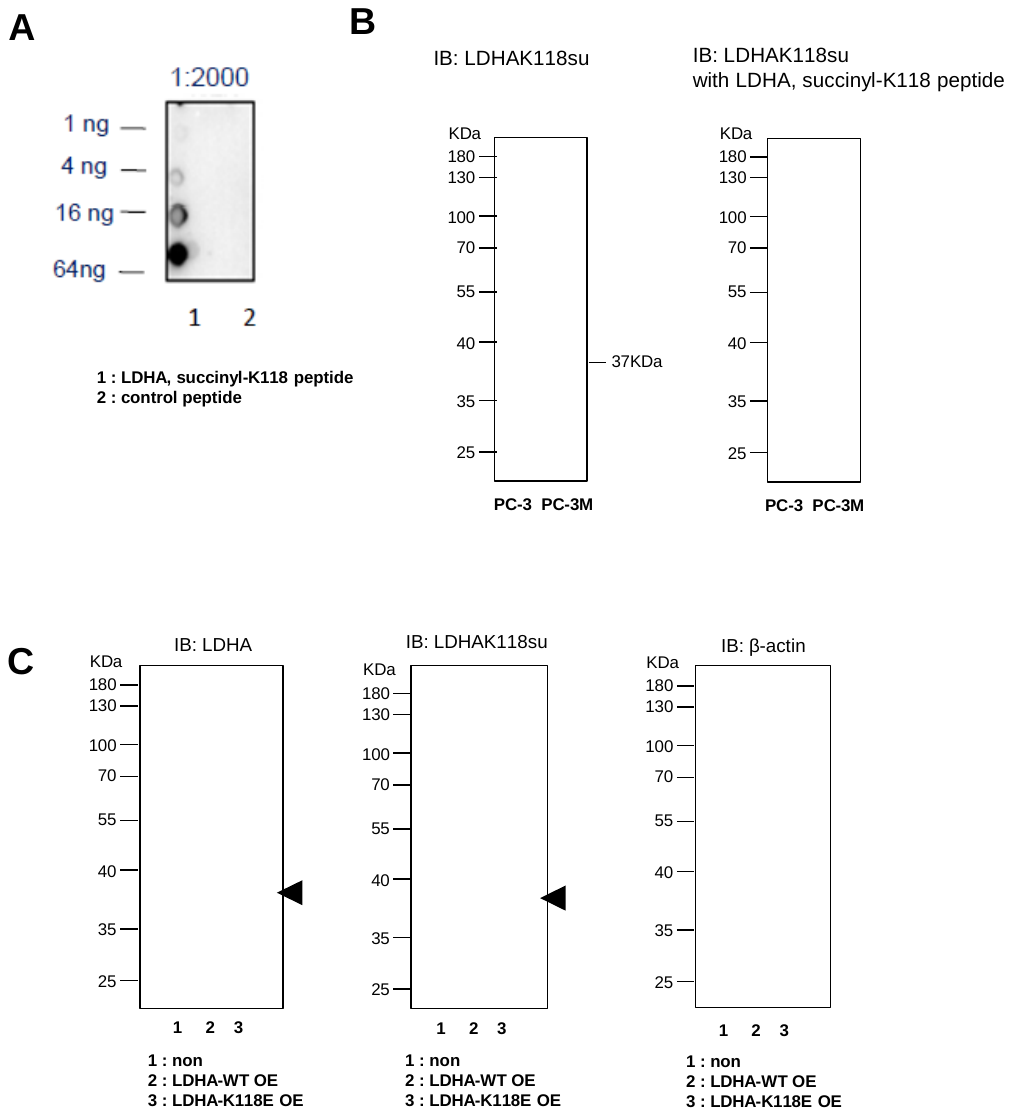

**Figure S7. Validation of selectivity of anti-LDHA-K118su**

**A**. Dot-blot assay showed the selectivity of anti-LDHA-K118su. Consistent with the ELISA result, the purified antibody showed very good immune-response with the modified peptides. The detection limit is lower than 4 ng. The signal doesn’t recognize the control peptide. **B**. Competitive assay of anti-LDHA-K118su in PC-3 and PC-3M cell lines using synthetic LDHA, succinyl-K118 peptides.
